# Behavioural plasticity in *Caenorhabditis elegans* navigating dynamic granular environments

**DOI:** 10.64898/2026.05.28.728193

**Authors:** Hongyi Xiao, Sima Maleki, Yuheng Pan, Adam Smith, Eleni Gourgou

**Affiliations:** Mechanical Engineering, University of Michigan, Ann Arbor, Michigan, United States; Biological Sciences, Wayne State University, Detroit, Michigan, United States; Computer Science & Engineering, University of Michigan, Ann Arbor, Michigan, United States

**Keywords:** *C. elegans*, behavioural plasticity, locomotion, microparticles, mechanosensation, touch sense, granular matter

## Abstract

The mechanisms that steer behavioural plasticity in complex environments remain poorly understood. Here, we introduce Peb2les, a quasi-two-dimensional granular arena, for the behavioural assessment of *Caenorhabditis elegans* nematodes, as they interact with deformable, dynamic terrains. Using customized deep-learning-based methods, we track both nematodes and particles to characterize coupled animal-environment dynamics. We find that nematodes’ locomotor performance depends on particle density and surface properties, and is largely independent of mechanosensation, whereas functional touch sense is required for the initial decision to enter rough-textured arenas. In addition, we identify a previously undescribed touch-seeking behaviour that partially depends on mechanosensation, in which *C. elegans* actively return to the granular arena after exiting. In parallel, locomoting nematodes rearrange and stack surrounding particles, continuously modifying the granular terrain. Together, these findings establish Peb2les as a versatile platform for probing the mechanism and limits of behavioural plasticity and locomotory adaptability of nematodes in sensory-enriched, soil-like environments, while also providing insight into the potential evolutionary origins of adaptive touch-seeking behaviours in animals.

## Introduction

Behavioural plasticity is of crucial importance for animals, especially when they experience novel and variable environments. It is remarkably sensitive to environmental changes [1] and it enables animals to respond to them in short time scales. However, the underlying neuronal mechanisms remain poorly understood. Such studies are extremely challenging in organisms with multi-million-neuron nervous systems. To this end, an animal that combines a compact and tractable nervous system with a rich behavioural repertoire is needed. *Caenorhabditis elegans* nematodes satisfy these criteria and allow for incisive genetic interventions and mechanistic insight [2, 3].

Indeed, *C. elegans* have been extensively used to map neuronal circuits that underpin a broad range of behaviours [4], including locomotion [5, 6], learning [7, 8], and decision making [9–12]. Despite targeted efforts [13–17], most studies investigate nematode behaviour on flat and largely featureless surfaces of agar-filled petri dishes. Such spaces do not reflect the qualities of *C. elegans’* natural habitat [18] (i.e. muddy soil, rotting plant material), which include granular matter with three-dimensional (3D) features. As a result, this model organism’s potential remains underexploited, especially for elucidating real-world behavioural plasticity [19] and therefore outlining the evolutionary origins of the trait [19].

We introduce a new type of nematode-friendly, quasi-two-dimensional (quasi-2D) granular arena, which allows nematodes to interact with microparticles dispersed on the surface of standard agar-based Nematode Growth Medium (NGM). We name these granular arenas *Peb2les* (<u>p</u>article-<u>e</u>nhanced <u>b</u>ehavioural quasi<u>2</u>D arenas for *C. e<u>le</u>gans*). Peb2les offer a sensory enriched environment, with varying particle density, texture, and physical properties. They are dynamic, because particle packing structures change as nematodes move through them.

*C. elegans* display a variety of mechanosensation-regulated behaviours [20–22]. The majority of previous work explores the behavioural outcome of touch input that is delivered in a localized, distributed manner, and results in animals moving away from gentle or harsh touches. However, it is not clear whether continuous and prolonged touch throughout the nematode body is perceived as aversive or attractive stimulus.

In Peb2le arenas, *C. elegans’* entire body is in almost constant touch with microparticles. We tracked the behaviour of wild type nematodes as they entered the arena, moved in the granular medium, exited, and re-entered, and we compared to that of mutants with impaired mechanosensation. We used customized deep learning-based algorithms to track the motion of both nematodes and particles. Our findings reveal a collection of dynamic behaviours that depend on particle properties and packing density, including nematodes returning into the sensory enriched arenas in a touch sense-dependent manner. We also characterize how nematodes induce particle motion and therefore alter the features of the granular packing structure, giving rise to bidirectional interactions between worms and their environment.

## Methods

### Preparation of Peb2le arenas and *C. elegans* nematodes

*C. elegans* nematodes were grown at 20°C, on standard Nematode Growth Medium (NGM) plates, with *E. coli* OP50 as food source [23]. The strains used were N2 wild type and CB1611 *mec-4(e1611)* (*Caenorhabditis* Genetics Center, University of Minnesota). Day 1 gravid adults were tested in two types of granular arenas: (i) polystyrene arenas, made of spherical, solid polystyrene particles (45 μm diameter, Polysciences, USA) with low surface energy (**1A, 1C**), (ii) glass arenas, made of spherical, hollow glass microbeads (60-70 μm diameter; Cospheric, USA) with high surface energy (**Fig. 1D, 3B**). For the assessment of entry attempts, diamond powder particles (60-80 μm diameter; Pureon, USA) with faceted shapes and low surface energy were used to create diamond arenas (**Fig. 4F**). Higher surface energy results in higher particle attraction, and consequently in particle clustering.

**Figure 1:**
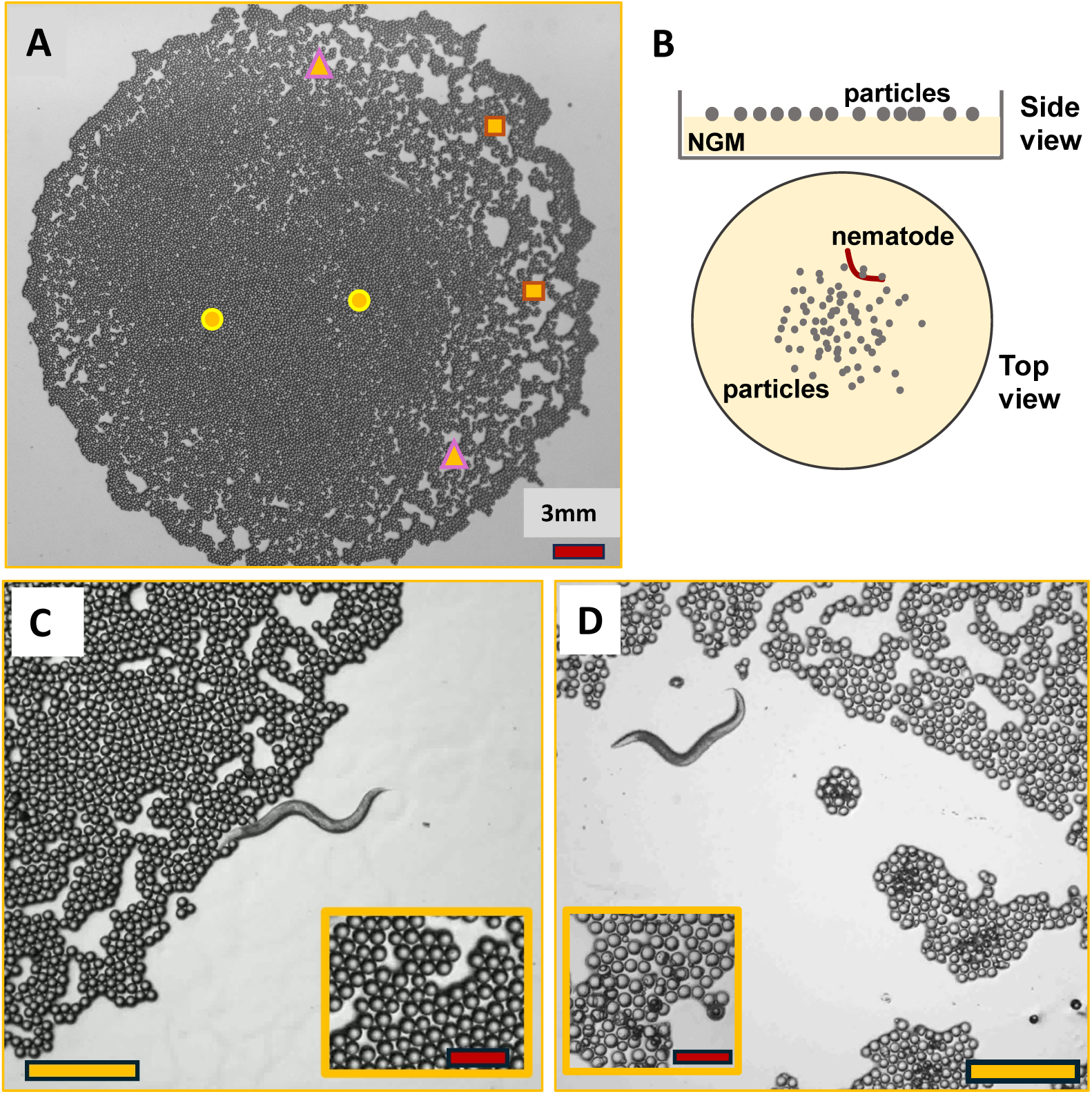
The granular terrain in Peb2le arenas. (A) Overview of a Peb2le arena (polystyrene) on an NGM plate. Symbols indicate areas of varying particle densities, circles: high density, triangles: medium density, squares: low density/no particles. Scale bar: 3 mm. (B) Schematic of NGM plate with Peb2le arena; side view (top), top view (bottom); particles not in scale with plate and worm. (C), (**D**) Particles used in Peb2le arenas, (**C**) polystyrene particles, (**D**) glass beads. Insets show magnified particle images. For particles sizes and properties, see Methods. Scale bars: 1 mm, scale bars of insets: 200 μm.

To create a Peb2le arena, particles suspended in M9 buffer [23], with a total volume of 150 μL, were dropped at the centre of a non-food seeded NGM plate, which remained in room temperature until excess moisture was absorbed. As a result, a layer of particles was formed at the centre of the plate, with the diameter of its perimeter around 1.5-1.7 cm, leaving the rest of the plate particle-free (**Fig. 1A, 1B**). The particles mostly formed a monolayer, and stacked particles were sparse. The spatial distribution of particles was heterogeneous, with tightly and loosely packed areas dispersed in the arena (**Fig. 1A**). Nematodes were introduced 5-10 mm outside the arena and were allowed to explore freely. In glass arenas worms have heavily impeded locomotion (see Results), so we also recorded videos with nematodes placed directly in the arena, to ensure we sufficiently capture their behaviour inside glass Peb2les. All videos were recorded with a DP22 camera and an Olympus SZ microscope, using CellSens imaging software (Olympus), with a frame rate of 13.74 frames per second.

### Nematode and particle tracking via Deep Learning

A nematode moving in a Peb2le arena is closely surrounded by particles, which appear as bright spots surrounded by dark halos and may overlap with the worm or other particles (**Fig. 2A, 2B**). Therefore, image processing-based identification of the worm and the particles, and acquisition of their temporal trajectories is challenging for conventional nematode or particle tracking algorithms. As a solution, we developed a deep learning-based segmentation method. For particle identification, we used the existing package Bellybutton [24], to identify all particle pixels in a given image. A similar algorithm was developed to identify pixels that belong to a worm. Based on a segmented image, we performed post-processing to extract the positions of the worm and the particles (**Fig. 2C**). For each worm, we assigned five body nodes (head, neck, midbody, hip, tail), defined at the 0^th^, 25^th^, 50^th^, 75^th^, and 100^th^ percentile of its total body length. A detailed description of the nematode and particle tracking methodology, body nodes definition, the steps taken to calculate displacement per second (**Fig. 3A** and **Fig. 4A**) and direction index *r* (**Fig. 3D** and **Fig. 4C**) are provided in the Appendix.

**Figure 2:**
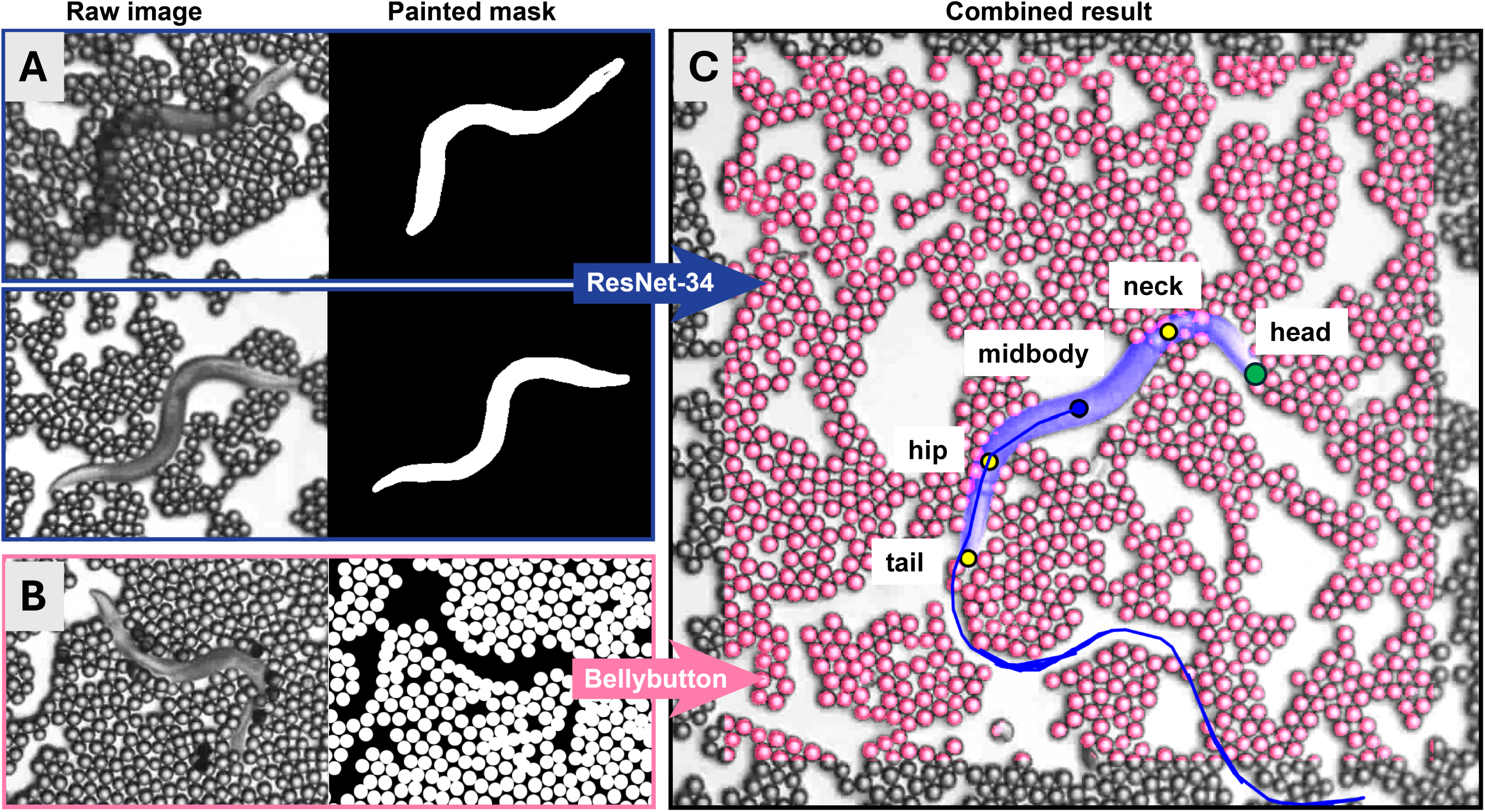
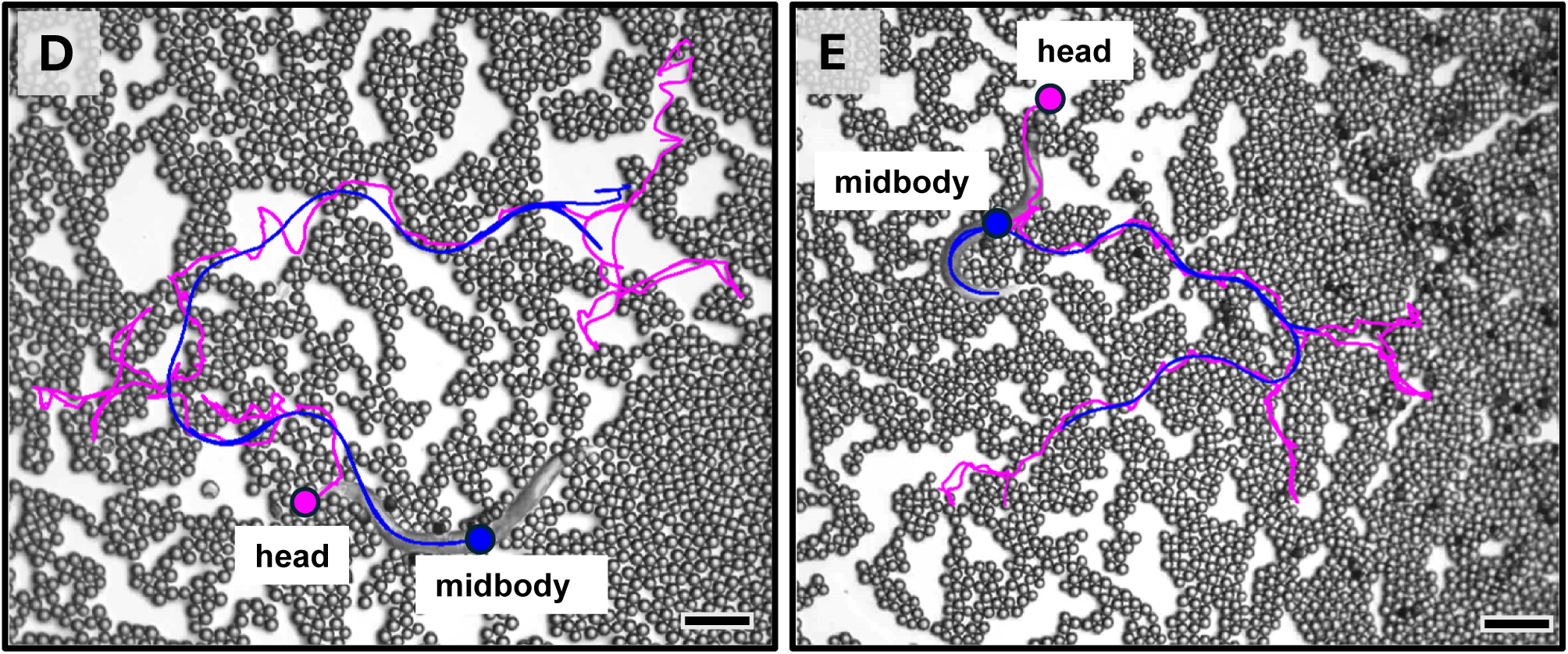
Image analysis for identification and tracking of nematodes and particles. (A) Examples of raw images and painted masks for worm tracking, using the ResNet-34 algorithm (see Methods and Appendix). (B) Example of raw image and mask for particle tracking, using the Bellybutton algorithm. (C) Combined tracking results with identified particle pixels, highlighted in pink, and worm pixels, highlighted in lavender. The identified worm body nodes are indicated with circles, green: head node, blue: midbody node, yellow: neck, hip, and tail nodes. Blue curve: trajectory of the midbody node. (D), (**E**) Examples of worm trajectories in polystyrene Peb2les, tracked over 1000 frames, *i.e.* ∼73 sec. Snapshots shown are the starting frame of each trajectory. Purple: head trajectories, blue: midbody trajectories. Scale bars: 350 μm. See Appendix for details on tracking methodology.

**Figure 3:**
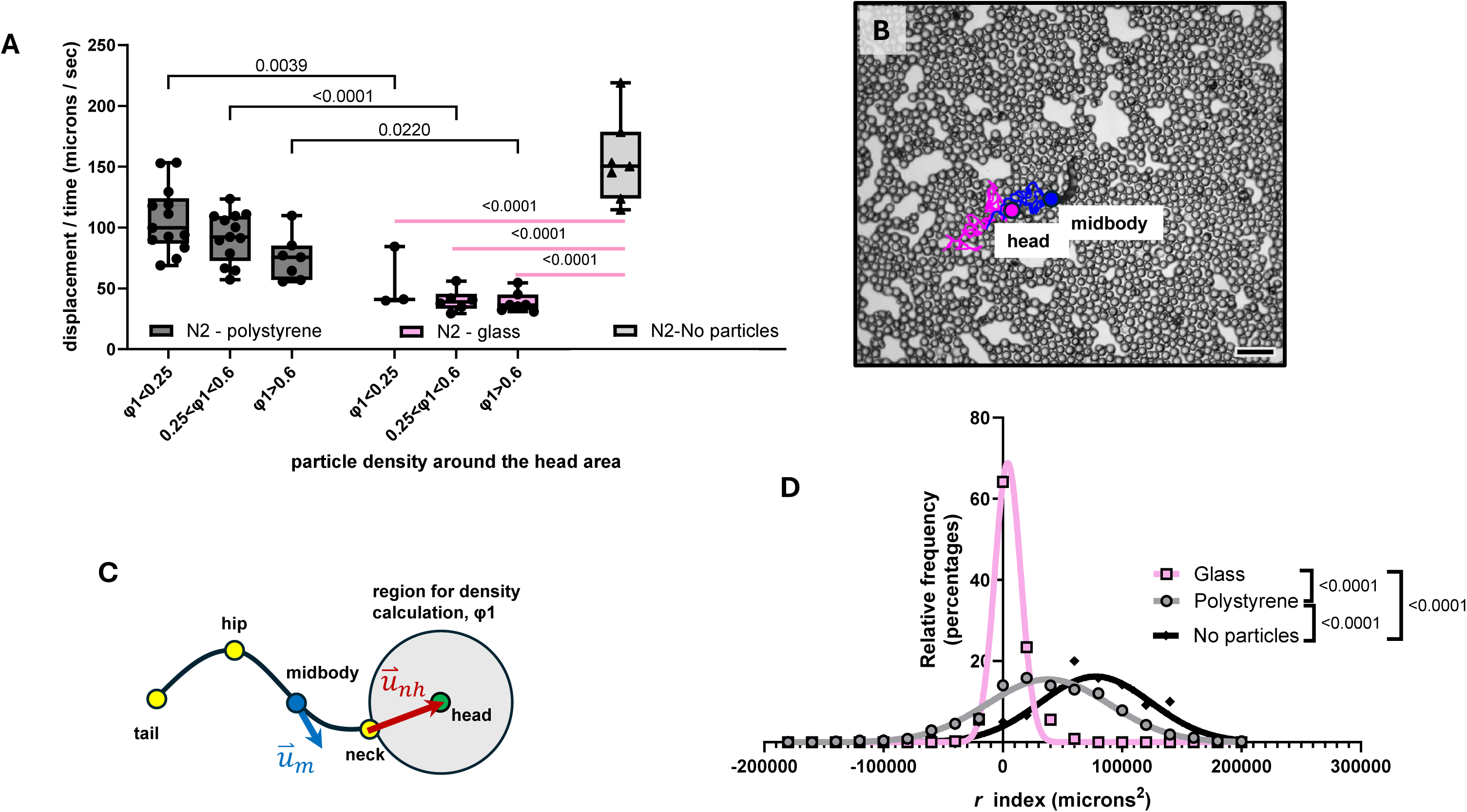
*C. elegans* locomotory performance depends on particle density and surface properties. **(A)** Displacement per second, of N2 wild type nematodes, in polystyrene (grey boxes) and glass (pink boxes) Peb2le arenas, in increasing particle density around the head area (*φ1*; see panel C and Fig. 1C; for calculation of particle density, see Appendix). Each dot represents displacement of one worm, n = 8-13 worms processed. The number of worms that visited areas of different densities varied. N2 wild type tracked in the absence of any particles (control) are also displayed (light grey). Horizontal lines: mean values, boxes: the central 50% of data, bars: min and max values. Brackets comparisons (black lines): two-way ANOVA, followed by Tukey’s multiple comparisons test; p-values between polystyrene and glass groups, for the same particle density. Straight (pink) lines comparisons: p-values between glass and control (no particles), one-way ANOVA followed by Tukey’s multiple comparisons test. Only statistically significant p-values (<0.05) are displayed. For comparisons between polystyrene N2 wild type nematodes and N2 control (no particles), see Fig. 3A. See also Supplementary Figure S2, for displacement with respect to particle density around other body areas. **(B)** Example of a worm’s midbody and head trajectories in a glass Peb2le, tracked over 1000 frames, *i.e.* ∼73 sec. Snapshot shown is the starting frame for both trajectories. Purple: head trajectory, blue: midbody trajectory. Scale bar: 350 μm. **(C)** Schematic of a skeletonized nematode, along with the five body nodes (see also Fig. 2C and Fig. S1A), showing the area around the head for the calculation of particle density φ1 (panel A) and vectors (u_m_: displacement vector; u_nh_: neck to head vector, see Appendix) that are used to calculate the direction index *r* (panel D). **(D)** Relative frequencies (percentages) distribution of direction index *r* for the assessment of backward/forward motion (see Methods and Appendix) of N2 wild type nematodes in polystyrene (grey) and glass (pink) arenas, as well as in the absence of any particles (control, black). Individual shapes indicate bin centre of histogram for frequency distribution; curves represent nonlinear fit for Gaussian distribution. *r*>0 indicates forward motion, *r*<0 indicates backward motion. See also Supplementary Figure S3 for data of individual worms for each group studied. One-way ANOVA followed by Tukey’s multiple comparisons test; p-values for comparisons between the means of the distributions provided in the legend.

### Nematode behaviour in high particle density, entry attempts, exits, and returns

In areas with densely packed particles, nematodes often crawled under the particles (**Fig. 4D**) or, in the case of glass beads, on top of them. For such events to be considered, nematodes had to be under or on top of particles with at least 1/3 of their body (**Fig. 4D**).

**Figure 4:**
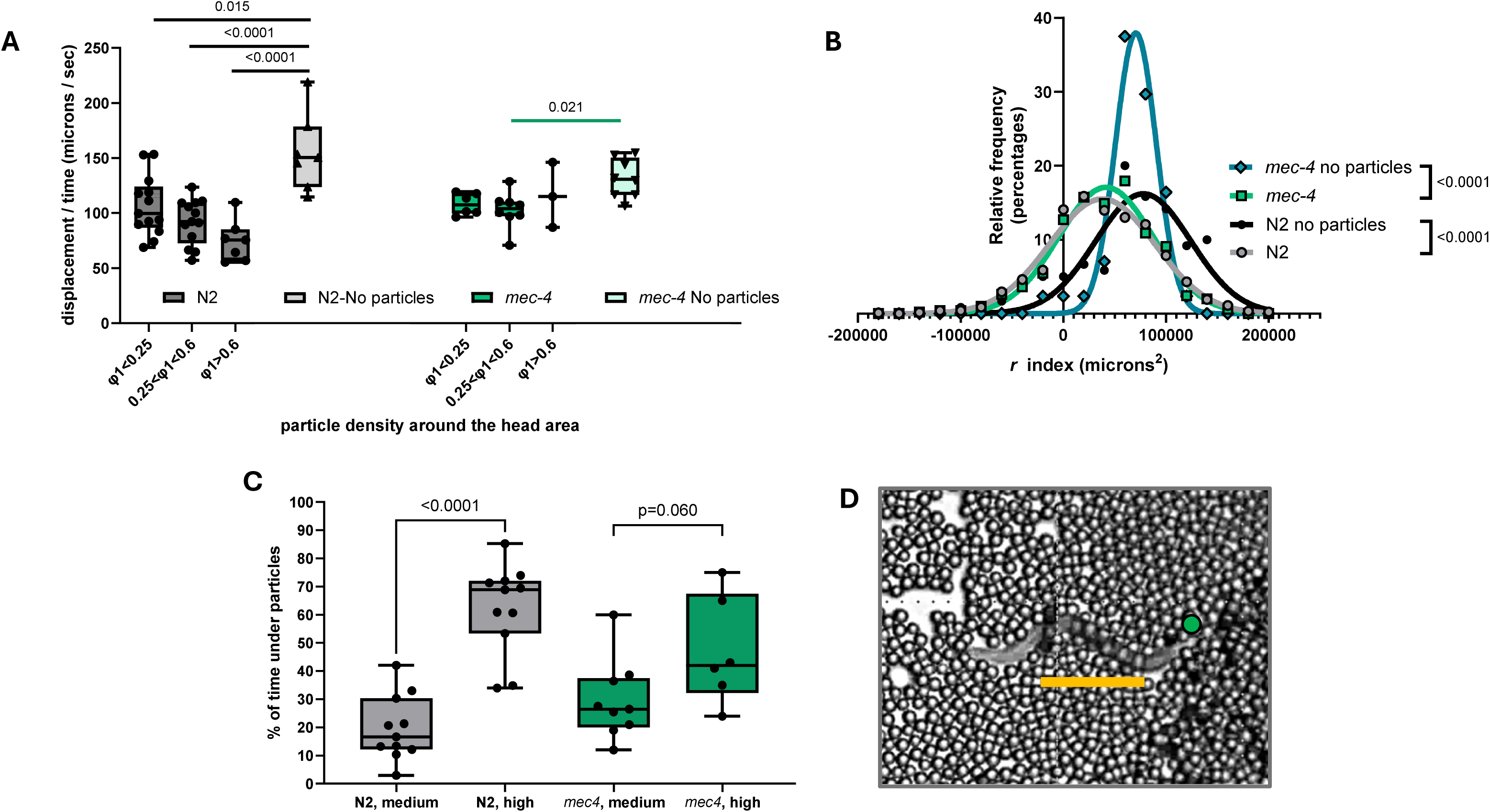
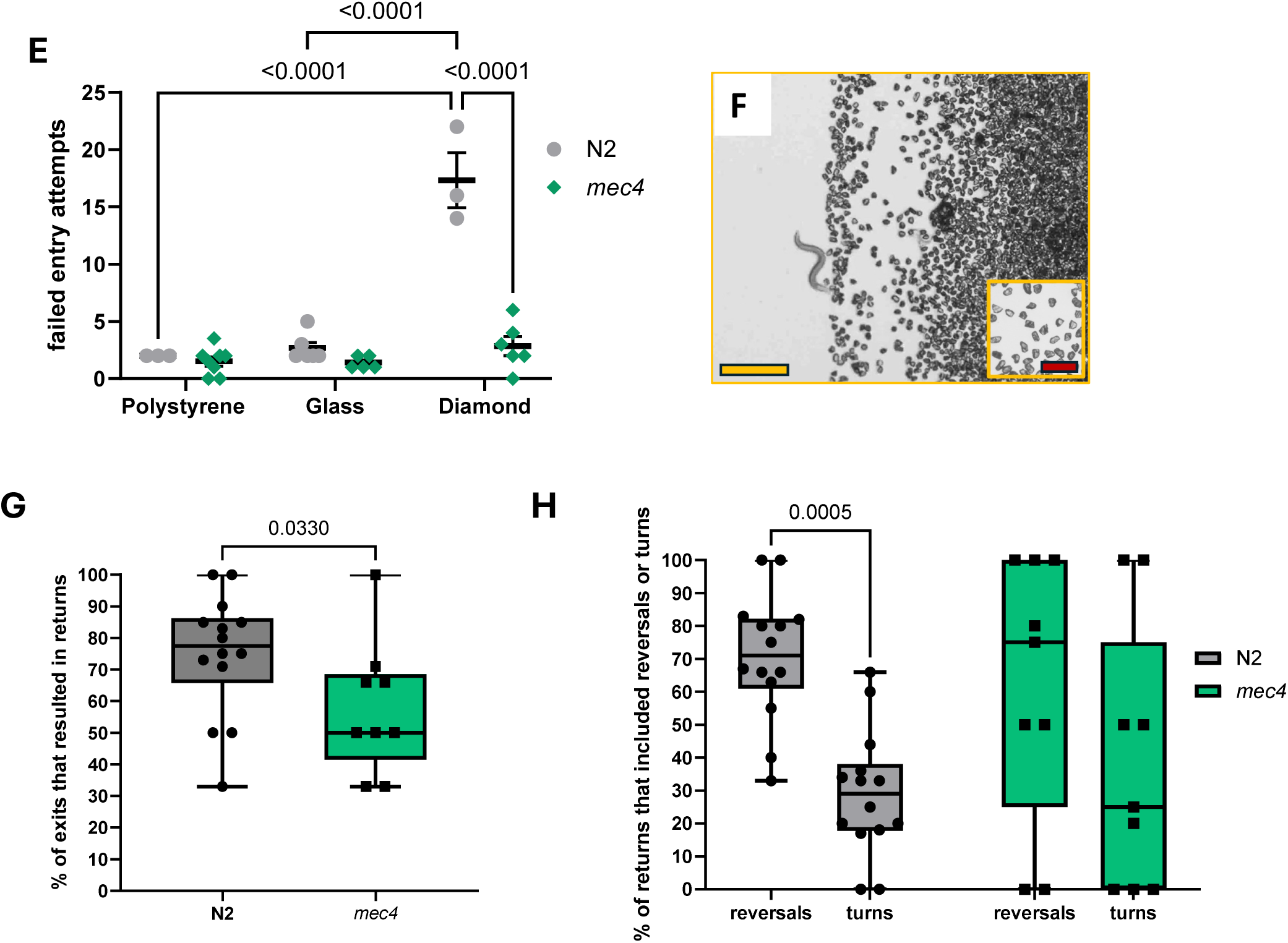
The role of *mec-4* in *C. elegans* locomotory performance, and touch-seeking behaviour in Peb2le arenas. **(A)** Displacement per second, of N2 wild type (grey boxes) and *mec-4(e1611)* nematodes (touch sense impaired; green boxes) in different densities of polystyrene Peb2le arenas. Each dot represents displacement of one worm, n = 6-13 worms processed for each strain. The number of worms that visited areas of different densities varied. N2 wild type (light grey) and *mec-4* (light green) tracked in the absence of any particles (controls) are also displayed. Straight lines comparisons (black and green): one-way ANOVA, followed by Tukey’s multiple comparisons test, between particle groups and the respective control (no particles). Comparisons between N2 and *mec-4* groups for the same particle density were performed by two-way ANOVA, followed by Tukey’s multiple comparisons test, and none of the comparisons yielded p-value<0.05. **(B)** Relative frequencies (percentages) distribution of direction index *r* for the assessment of backward/forward motion (see Methods and Appendix) of N2 wild type (grey) and *mec-4* nematodes (green) in polystyrene arenas, as well as in the absence of any particles (N2 control, black; *mec-4* control, dark turquoise). Individual shapes indicate bin centre of histogram for frequency distribution; curves represent nonlinear fit for Gaussian distribution. *r*>0 indicates forward motion, *r*<0 indicates backward motion. One-way ANOVA followed by Tukey’s multiple comparisons test; p-values for comparisons between the means of the distributions provided in the legend. (C) Left: % of time spent under polystyrene particles, for N2 wild type (grey) and *mec-4(e1611)* (green) nematodes, depending on particle density (medium: 0.3< ¢_c_<0.5, high: ¢_c_ >0.5). Note that particle density in this case is defined with respect to the area in front of the worm (Supplementary Fig. S2B). Two-way ANOVA, followed by Fisher’s multiple comparisons test. (D) A nematode crawling partially through and partially under (yellow line) polystyrene particles, head indicated with a green dot. (E) Wild type N2 and *mec-4(e1611)* nematodes’ failed attempts before they finally enter polystyrene, glass, and diamond Peb2le arenas (for event definitions see Methods). Each dot represents entrance attempts of one nematode. Lines: mean values, error bars: SEM. Two-way ANOVA, followed by Tukey’s multiple comparisons test. (F) Diamond powder Peb2le arena, scale bar: 1 mm, scale bar of inset: 200 μm. (G) % of exits that resulted in returns (for event definitions see Methods), for N2 wild type (grey) and *mec-4(e1611)* (green) nematodes. Mann Whitney test. See also Supplementary Fig. S4. (H) % of returns that were realized by either a reversal or a turn (for event definitions see Methods), for N2 wild type (grey) and *mec-4(e1611)* (green) nematodes. Wilcoxon matched-pairs signed rank test. All panels: only p-values <0.05 included; horizontal lines: mean values, boxes: the central 50% of data, bars: min and max values; p-values on graphs, as described.

Failed entry attempts (**Fig. 4E**) were considered events in which a forward-moving worm’s nose tip touched the particles of the outer arena perimeter, immediately followed by a reversal. Successful entry attempts were considered events in which a forward-moving worm’s nose touched the particles of the outer perimeter, followed by forward motion for at least two undulation cycles.

Exits (**Fig. 4G**) were consider events in which a nematode’s head and the anterior part of the body emerged out of the arena, regardless of whether the rest of the body followed. Returns (**Fig. 4G, 4H**) were defined as events in which nematodes turned back into the arena in less than 40 sec after exit. Returns were further distinguished into reversals and turns (**Fig. 4H, Fig. S4**). For the former, nematodes performed a reversal back into the arena. For the latter, nematodes performed a full pirouette, an omega turn or turned back into the arena via a sharp turn. Entry and exit events were counted regardless of whether worms crawled through or under particles during their exit or upon entry.

### Chemotaxis assay

The chemotaxis assay (**Fig. 5A**) was performed according to a well-established protocol [25]. A synchronized population of Day 1 adult hermaphrodites was obtained, following standard process [23]. A four-quadrant assay NGM plate was prepared [25], test and control quadrants were alternated across experiments, and the chemotaxis index was calculated as *Chemotaxis Index*= (N_test_ − N_control_)/(N_total_), where N_test_ is the number of worms in both test quadrants, N_control_ is the number of worms in both control quadrants, and N_total_ is the number of total scored worms. Five to eight replicates for each condition were performed for each assay, across different days. Details are provided in the Appendix.

**Figure 5:**
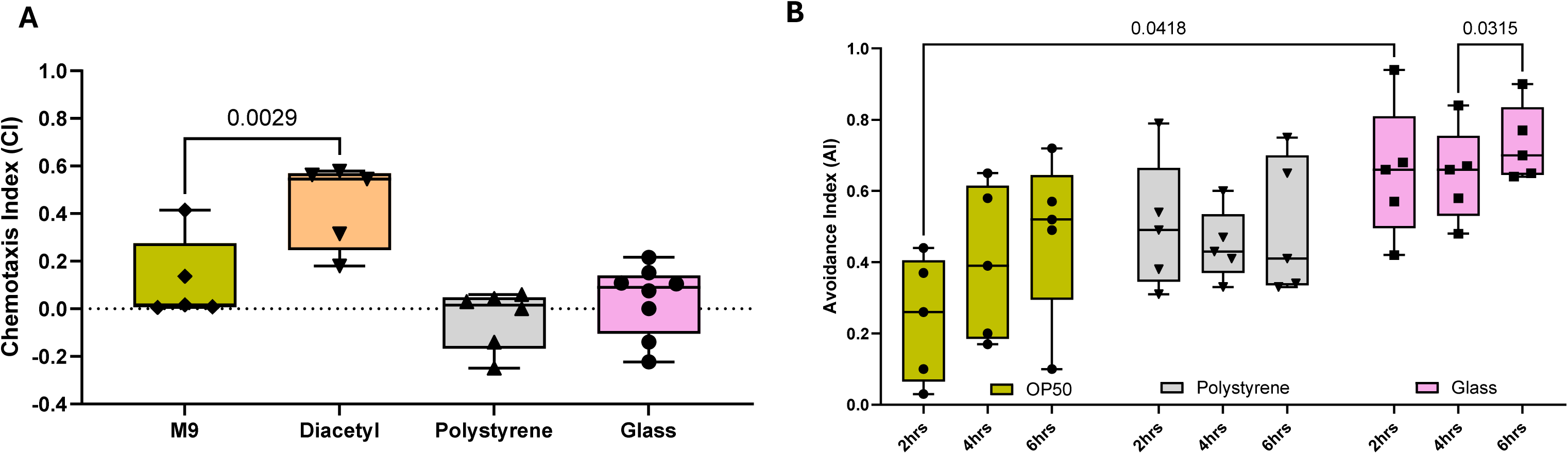
Short term chemotaxis for Peb2le-forming particles and long term avoidance of Peb2le arenas. (A) Chemotaxis index for N2 wild type nematodes and particles used in Peb2le arenas, over 1 hr. Particles were mixed with M9, as was the case in experiments of Figs. 3 and 4. Comparisons against M9 buffer (carrier buffer, control). Diacetyl provided for reference of positive chemotaxis. One-way ANOVA, followed by Tukey’s multiple comparisons test. Shapes indicate n = 5-8 independent assays for each group tested. Only p-values<0.05 are shown. (B) Avoidance index at different time points (2 hr, 4 hr, 6 hr) for N2 wild type nematodes and the particles used in Peb2le arenas. Particles were mixed with *E. coli* OP50. Avoidance index for plain OP50 bacterial lawn, no particles, is provided for comparison. Two-way ANOVA, followed by Tukey’s multiple comparisons test. Shapes indicate n = 5 independent assays for each group tested. Only p-values<0.05 are shown. For avoidance index when particles are mixed with M9 buffer, see Supplementary Fig. S5.

### Avoidance assay

For the avoidance assay (**Fig. 5B**), a previously published protocol was followed [26]. A synchronized population of Day 1 adult hermaphrodites was obtained, following standard process [23]. Results were quantified by calculating the fraction of worms staying on the particle/bacterial lawn over the total number of worms in the assay plate, as *Avoidance Index* = N_off_/N_total_, where N_off_ is the number of worms outside the lawn and N_total_ is the number of total worms on the plate. Five replicates for each condition were performed for each assay, across different days. Details are provided in the Appendix.

### Particle packing analysis

To elucidate the interplay between a crawling worm and its granular environment, we characterized the particle kinematics and packing structure. This required identifying each individual particle’s centre location. We analysed videos of worms in polystyrene arenas, where particles form primarily monolayers. We could track most of them, except for the rare, stacked ones (**Fig. 6A**, white arrow). We calculated the particle velocity, v_p_, based on the consecutive positions of a particle over an interval of 0.73 s, corresponding to 10 video frames.

**Figure 6:**
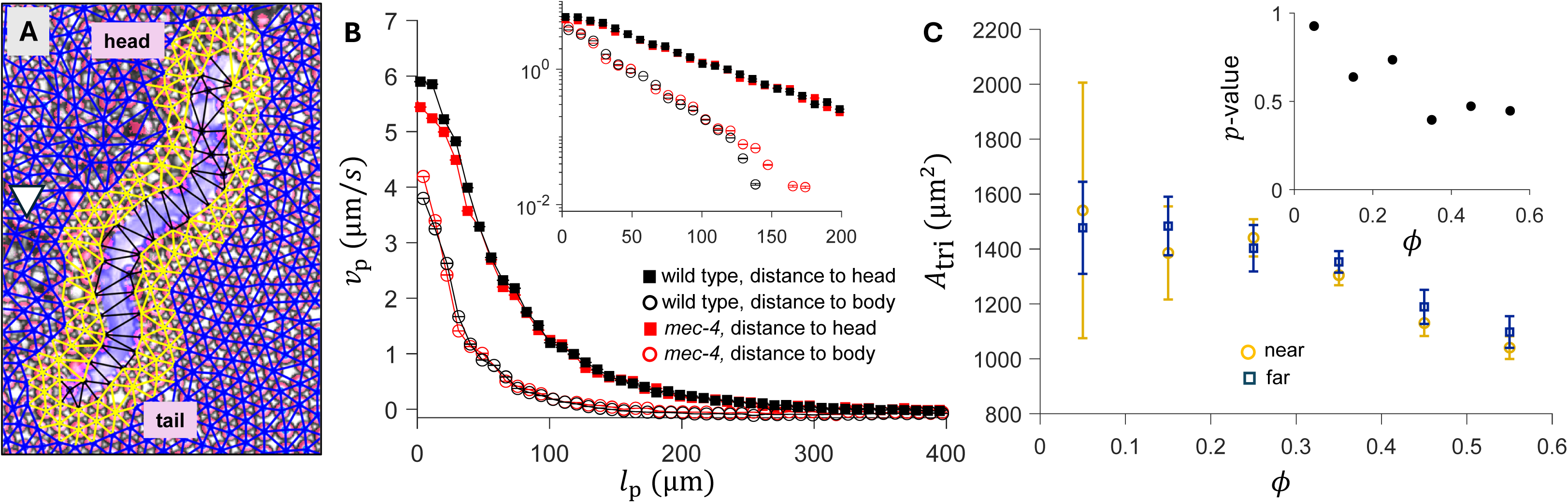
Particle-based analysis. (A) Visualization of tracked particle centers (blue and yellow dots) and Delaunay triangulation (blue and yellow lines, see Appendix). Yellow dots and lines: particles in the proximity of the worm body; blue dots and lines: particles far from the worm body. White arrow points at stacked particles, while most of the particles form a monolayer. (B) Particle velocity v_p_ as a function of the particle-to-worm distance l_p_ for wild type (black) and *mec-4* (red) nematodes. Solid symbols: particle velocity *vs*. distance to the head node, hollow symbols: particle velocity *vs*. shortest distance to the worm’s body contour. Inset: same data, plotted on a log-lin scale. Analysis performed on N2 wild type nematodes. (C) Bin-averaged triangle area A_tri_ with respect to packing density ∅. Orange circles: triangles near the worm; dark blue squares: triangles far from the worm, as visualized in (A). Error bars: standard errors measured from five trials having different initial particle arrangements. Inset: Student’s t-test, p-values calculated based on the near-field and far-field triangles at each packing density. See Appendix for details on methodology.

To determine how a moving nematode rearranges nearby particles, we characterized the particle packing structure (**Fig. 6A**). We identified particle centres and performed Delaunay triangulation, a common and effective method for 2D and quasi-2D granular systems [27–29].

To detect nematode-induced broad range modifications of the Peb2le arena we compared images of the entire arena taken before and 3 hr after worms were introduced and roamed in the arena (**Fig. 7A**). A description of the methodology, including a Fast Fourier Transform analysis, (**Fig. 7E**) is provided in the Appendix.

**Figure 7:**
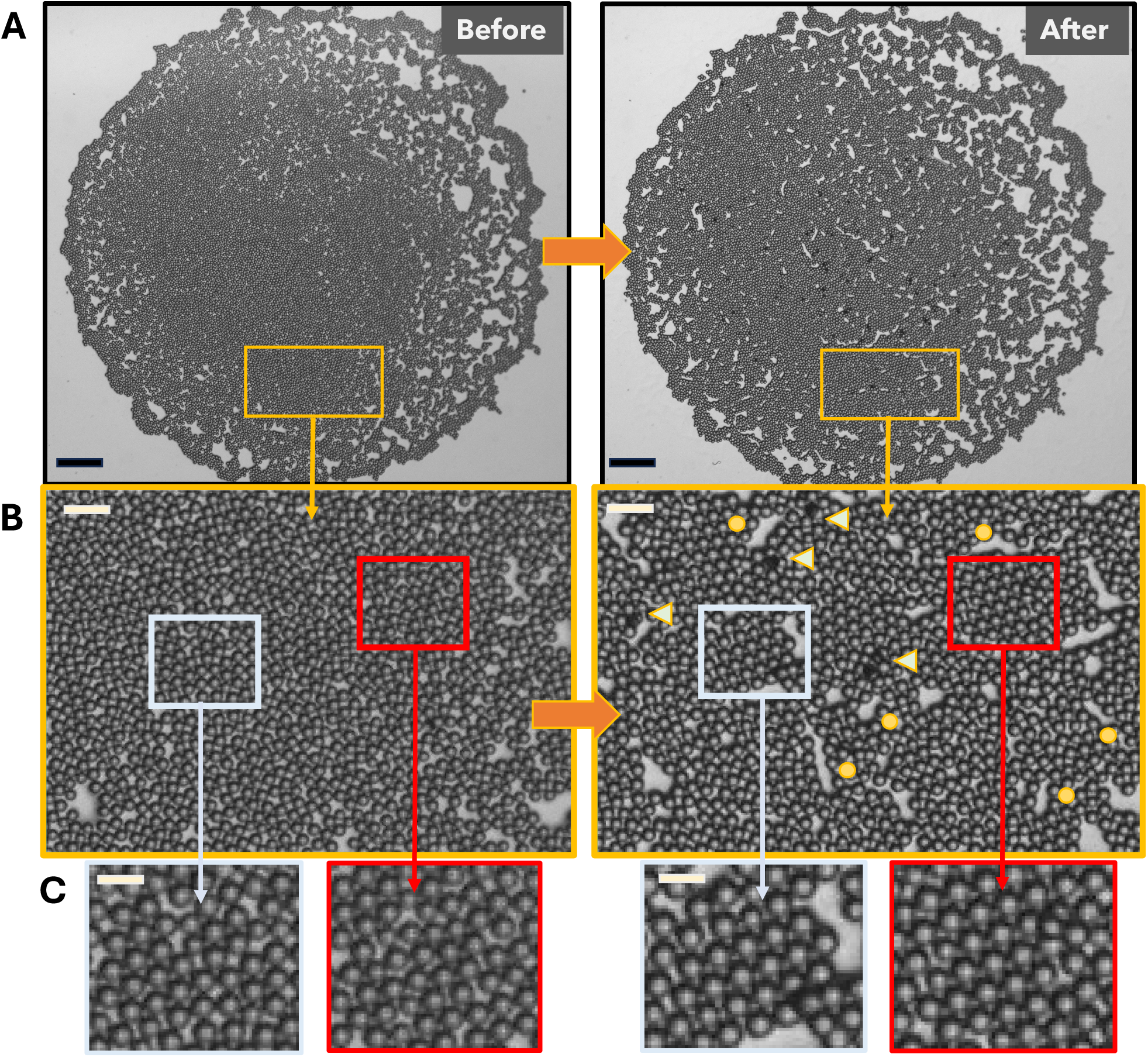
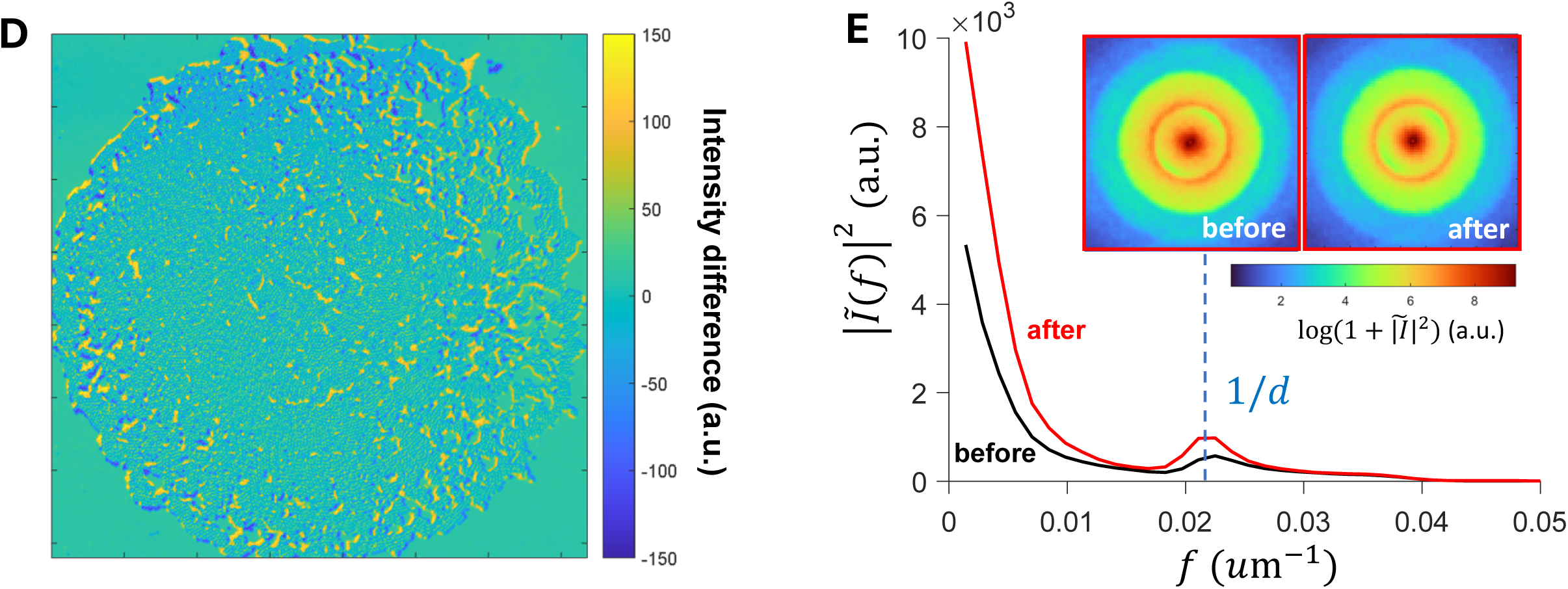
Locomoting nematodes reshape the granular arena. (A) A polystyrene Peb2le arena before (left) and after (right) three nematodes roamed freely for ∼3 hr, scale bars: 2.5 mm. (B) Images in yellow frames are higher magnifications of square arena regions with the same color frame as in A panels. Right (after): yellow circles are placed next to newly shaped void areas, and arrows point at new particle stacks; not all such formations are labeled. Scale bars: 250 μm. (C) Images in light blue and crimson frames are higher magnifications of square arena regions with the same color frame as in B panels. Random packings of particles (left two panels) are converted into organized, crystal-like packings after nematodes traveled through (right two panels). Scale bars: 100 μm. (D) Difference in intensity (arbitrary units) between before and after images of panel A. (E) Radially-averaged intensity, |*Ĩ*(f)|^2^ *vs*. wave vector, f, of two-dimensional Fourier transform for the before (black) and after (red) images of (A), arbitrary units. Insets: Fourier transform results for the before/after images.

### Statistical analysis

Each experiment was conducted over a period of 2-4 weeks. Comparisons between nematode groups were performed as described in figure captions. All statistical tests were performed using GraphPad Prism 11 (GraphPad, USA), unless stated otherwise. In all cases, comparisons were considered statistically significant when *p*-value<0.05.

## Results

### *C. elegans* sinusoidal crawling motion is maintained in the polystyrene Peb2le arena

When crawling on an NGM plate, *C. elegans* locomotion presents characteristic dorsoventral undulations [30, 31]. As a result, the nematode follows a sinusoidal path. Locomotion is realized by forward (head-first) and reverse (tail-first) crawling, occasionally interrupted by pirouettes [32] or omega turns [33]. In polystyrene Peb2les (**Fig. 1A, 1C**), *C. elegans* crawled on NGM, surrounded by particles, packed in varying densities (**Fig. 1, 2A, and Supplementary Videos V1-V2**). Crawling was performed through undulatory motion, with clear dorsoventral bends, executing a continuous sinusoidal-like wave (**Figs. 2C, 2D, 2E**, blue trajectory; and **Supplementary Video V3**). Worms crawled mostly forward, and they occasionally engaged in reversals, both captured successfully by the tracking algorithm (**Supplementary Video V3**), which also captured side-to-side head sweeps, linked to *C. elegans’* foraging behaviour [31] (**Fig. 2D, 2E**, pink trajectory). Omega turns and pirouettes were seldom performed (**Supplementary Videos V1-V2**). Overall, locomotion in polystyrene Peb2les maintains most of the hallmark features of standard *C. elegans* crawling behaviour but complex manoeuvres, like pirouettes, are rare. This suggests that *C. elegans* adapt the palette of locomotory expressions depending on the environment’s physical properties.

### Locomotory behaviour depends on particle density and surface properties

We tracked nematode locomotion in varying particle densities and compared it with locomotion in the absence of particles. When traversing a polystyrene Peb2le, nematodes’ mean displacement per time unit is reduced, compared to particle-free arenas (**Fig. 3A**). Next, we asked whether there is a correlation between a worm’s displacement per time unit and its surrounding environment. To test this, we measured particle packing density (i.e., particle area fraction) around five body areas, each centred around a body node (**Fig. 2C**). Displacement per time unit decreases with increasing particle density around the first body node, *i.e*., the head area (**Fig. 3A** and **Fig. 3C**, *φ_1_*, grey circle; for calculation of particle density around body nodes see Appendix and **Fig. S1A**) and around the second body node, *i.e*., the neck area (*φ_2_*, **Fig. S2**). There is no correlation with particle density around the remaining three body nodes at the mid and posterior body (*φ_3_*-*φ_5_*, **Fig. S2**). Therefore, density of granular medium around the anterior body (head and neck areas) is important for adjusting locomotory speed, while density around the midbody and posterior area is less salient. Moreover, the mean of the cumulative frequency distribution of the *r* index (direction index for the assessment of forward/backward motion, see Methods and Appendix) decreases when *C. elegans* move in Peb2les, compared to particle-free arenas (**Fig. 3D**, grey and black curve, respectively, and **Fig. S3**). Nematodes in the presence of polystyrene particles move backward more, compared to particle-free arenas, however they maintain an overall forward motion.

Next, we asked whether surface properties of granules affect the performance of crawling nematodes. To test this, we used glass beads, which are spherical (**Fig. 1D**) like polystyrene particles, but have higher wettability and surface energy [34], resulting in them forming clusters. In glass Peb2les, nematodes have a hard time locomoting (**Supplementary Videos V4-V5**). Their displacement per second is significantly decreased (**Fig. 3A**), compared to that in both polystyrene and particle-free arenas. Although nematodes can easily push around individual glass beads (**Supplementary Video V6**), they cannot break through clumped glass particles. Therefore, most (∼50-70%) of the time they climb on top of the glass beads, where their undulatory motion does not effectively generate thrust. As a result, they move forward very slowly (**Fig. 3A**). Combined with the condensed trajectory captured by the tracking algorithm (**Fig. 3B**) and the cumulative frequency distribution of *r* index that has a near-zero mean (**Fig. 3D**, pink line, and **Fig. S3**), these findings indicate that nematode locomotion in glass Peb2les is severely obstructed, as *C. elegans* remain mostly stationary.

### Impaired touch sense does not affect nematodes’ locomotory behaviour in Peb2les

We asked whether touch sense provides sensory cues that are important for locomotory adaptability in Peb2les. To this end, we tested *mec-4(e1611)* nematodes, which have impaired touch sense [35, 36]. We found that, like wild type nematodes, *mec-4* worms move more slowly in the presence of polystyrene particles than in particle-free arenas (**Fig. 4A**). Displacement per second of *mec-4* is not different compared to that of their wild type counterparts (**Fig. 4A**). At the same time, it seems not to be significantly affected by particle density, although the number of worms detected in high density areas is smaller (fewer data for φ1>0.6, **Fig. 4A**, green). Moreover, the mean of the frequency distribution of r index is not significantly different between *mec-4* and wild type in polystyrene arenas (**Fig. 4B**, grey and green curves), nor between *mec-4* and N2 in the absence of particles (**Fig. 4B**, black and dark green curves). However, the r index of *mec-4* conforms more around the mean of the distribution (**Fig. 4B**, dark green), indicating less variance among mutant worms.

### Nematodes crawl under the particles in dense areas of polystyrene Peb2les

In areas of the polystyrene arenas with medium (0.3<*φ_c_*<0.5) and high (*φ_c_*>0.5) particle density (**Supplementary Figure S1B)**, nematodes often crawl under the particles, instead of pushing through them (**Fig. 4C, 4D, Supplementary Videos V1-V2**). The fraction of time spent crawling under the particles increases significantly in highly dense areas *vs* medium dense (68% *vs* 18%, **Fig. 4C**, grey). This behaviour is observed also in *mec-4(e1611)* worms with impaired mechanosensation, even though the difference between the mean values at high and medium densities is smaller (40% *vs* 25%, **Fig. 4C**, green), N2 and *mec-4* nematodes behave similarly in medium densities (*p*-value>0.05, **Fig. 4C**). When comparing the behaviour of the two strains in high densities, the difference is only borderline non-significant (*p*-value=0.053, **Fig. 4C**), with N2 worms spending 68% and *mec-4* worms spending only 40% of their time crawling under the particles. Therefore, the switch to crawling under the particles depends on particle density, and touch sense is not found to be the primary sensory modality that informs this behaviour. At low particle density (*φ_c_*<0.3), this behaviour is not observed for either strain (0% of time spent under the particles, data not shown).

### Decision to enter the arena depends on particle properties and is mediated by touch sense

We analysed nematodes’ first encounter with the particles at the perimeter of the arena. N2 wild type enter polystyrene and glass arenas after a similar, low number (0-5 in total) of failed attempts (**Fig. 4E**, grey), indicating a comfortable initial interaction with either particle type. The same is true for *mec-4* nematodes (**Fig. 4E**, green), suggesting that impaired touch sense does not affect the outcome of entry attempts in either type of Peb2le arena. Note that in the case of glass arenas, the difficulty for *C. elegans* is to locomote after their head and neck have crossed the outer perimeter, that is after the animal has made the decision to enter (**Supplementary Video V6**).

Both polystyrene and glass arenas are made of spherical particles (**Fig. 1C, 1D**). We asked whether an entry is similarly easy when the overall medium roughness increases. To this end, we used diamond powder that features irregular-shaped, multifaceted particles (**Fig. 4F**). In this case, N2 wild type nematodes execute more failed attempts (13-22 in total) before they finally enter (**Fig. 4E**, grey, and **Supplementary Videos V7-V8**), while *mec-4* worms require significantly fewer (0-7 attempts, **Fig. 4E**, grey, and **Supplementary Videos V9-V10**). These findings suggest that *mec-4* worms do not distinguish between spherical and multifaceted particles as they decide to enter a particle area. Moreover, they suggest that nematodes perceive certain textures as less inviting than others and this affects their decision to enter a granular arena.

### Nematodes return to the polystyrene Peb2le

Nematodes that exit the polystyrene arena are very likely to re-enter, as 78% of N2 wild type exits are followed by a return (**Fig. 4G**, grey). Out of these return events, the majority, *i.e.* 71%, are realized via a reversal (**Fig. S4**, and **Supplementary Video V1**, 0:35-0:38) and the remaining 29% via pirouettes, omega turns or sharp turns (**Fig. 4H**, grey, **Fig. S4**, and **Supplementary Video V2**, 0:08-0:22). With respect to *mec-4* nematodes, only half of the exits (50%) are followed by a return (**Fig. 4G**, green). Out of these, 73% are realized via a reversal, and 27% are performed via other types of turns (**Fig. 4H**, green, and **Fig. S4**), in a ratio similar to that of N2 worms. Therefore, returning behaviour is significantly reduced when touch sense is impaired, while the returning manoeuvres remain the same.

It is noted that most nematodes (∼95%) exit the arena after spending more than 15 min in it. The few (<5%) that exit the arena quickly, *i.e.* in less than 3 min after their first entry, are less likely to return, and they have not been included in the above analysis. This applies to both N2 and *mec-4* worms.

### *C. elegans* do not sense particles through chemical cues

To test whether nematodes’ behaviour is affected by chemical cues released from the particles, we performed a chemotaxis assay [25]. The chemotaxis index for all particles was similar to the one of the control (**Fig. 5A**). Therefore, particles present no chemical cues that attract or repel nematodes.

We also asked whether *C. elegans* avoid Peb2les over longer periods of time. To test this, we performed an avoidance assay [26] for 2 hr, 4 hr, and 6 hr. The avoidance index for both polystyrene and glass arenas, where particles were mixed with OP50, was not significantly different than the avoidance index for plain OP50 (**Fig. 5B**), with the exception of glass particles at the 2-hr time point. Therefore, nematodes do not avoid polystyrene particles and steer away from glass beads. When either particle type was mixed with M9 instead of OP50, the avoidance index was again similar to the one of the control, namely plain M9 (**Fig. S5**). In this case, the higher avoidance index for all cases, including the control, can be attributed to *C. elegans* looking for food.

### Particle kinematics and rearrangement by traveling nematodes

As nematodes moved through the polystyrene Peb2les, they pushed and relocated particles. To quantify this, we performed particle kinematics analysis, examining the particle velocity, v_p_, as a function of the particle distance from a worm, l_p_, which was defined as either distance from the worm’s body or distance from the head node (**Fig. 6A**, see Appendix). For both N2 wild type and *mec-4* mutants, particle motion decays exponentially with increasing distance, as shown when results are plotted in a semi-log scale (**Fig. 6B**). Particles closer to the head display larger worm-induced motion than particles elsewhere and have larger decay length.

To determine how moving nematodes change the arrangement of adjacent particles, we drew Delaunay triangles (**Supplementary Video V3**, and Appendix) and categorised them into near-field (yellow) and far-field (blue) triangles (**Fig. 6A**). The difference between near- and far-field triangles was statistically insignificant at all particle densities (*p*-values >0.05, **Fig. 6C**).

Nematodes reshaped the arena landscape more broadly, over a longer period of time (**Fig. 7A**, before /after). New void areas were created (circles in **Fig. 7B**, right *vs* left panel) and particles were stacked (darker spots, triangles in **Fig. 7B**, right *vs* left panel), often next to the new void areas. In addition, particles in several regions were packed in a more ordered manner (**Fig. 7C**, right *vs* left panel). These changes were revealed via the image intensity differencing result (**Fig. 7D**). The Fourier transform-based quantification of the sub-domains with significant packing structure difference (**Fig. 7E**, and Appendix) confirms and highlights further changes in particle packing and in the ratio of void versus particle-covered areas.

## Discussion

### Peb2les: a new granular arena for the assessment of *C. elegans* behaviour

*C. elegans* nematodes’ natural habitat [37, 38], *i.e*., muddy soil and decomposing plant material, often consists, in part or in whole, of granular matter. In a lab setting, according to the gold standard, *C. elegans* are cultured on the flat surface of petri dishes filled with agar-based NGM [23]. Several attempts have been made to create more realistic experimental terrains [39, 40], including 3D-printed physical barriers made of NGM [41], dehydrated fruit-based scaffolds [40], and cotton gauze [42]. These environments are an advancement compared to the standard NGM dishes, but they often have limited reproducibility and compatibility with imaging techniques or require sophisticated fabrication methods. Soil-like granularity has been introduced in microfluidic devices with bumps or microposts that imitate dirt particles [43]. These microstructures are stationary, their texture cannot be modified unless the device is remade, and the environment does not change as nematodes move in it. Therefore, the mechanism of behavioural plasticity in a dynamic environment cannot be adequately investigated, and dynamic body-environment interactions cannot be explored.

*C. elegans* have been reported to move along sand columns, in a way that depends on particle size and levels of moisture [44]. Granularity of *C. elegans* habitat has also been introduced in depositions of soft dextran gel Sephadex beads, that successfully mimic bacterial lawns [45–47]. Sephadex gel beads are in the range of 20-50 μm [45], and they form a continuous matrix [47], rather than a layer of stand-alone granules, as is the case with microparticles in Peb2les (**Fig. 1**). Recent work assesses *C. elegans* locomotive behaviour in 3D granular matrices, made out of soft hydrogel beads [48], which constitute a nematode-friendly environment that can be elastically deformed and reversibly displaced by the nematodes’ motion.

Peb2les (**Fig. 1**) are deformable, dynamic, and quasi-2D arenas, *i.e.,* 2D enriched with 3D elements. They resemble soil granules more closely, because they include particles that are stiff and frictional, and their size is close to the range of silt granules [49]. Moreover, void areas in Peb2les resemble pockets of air found in soil [50]. Peb2les are created by spreading microparticles on the surface of NGM dishes, therefore nematodes still crawl on their familiar agar surface and display their well-characterized locomotory behaviour. At the same time, sensory input is enhanced, compared to the featureless, flat-surfaced standard NGM dishes. Therefore, Peb2les maintain the advantages of NGM plates, and in addition they allow for imaging and tracking (**Fig. 2**) of nematodes in a dynamic, sensory enriched terrain. These attributes make Peb2les more realistic and more informative than most of the commonly used behavioural platforms for nematodes.

Peb2les are a versatile type of behavioural arena that can be made with a variety of particles, in our case polystyrene, glass, and diamond (**Fig. 1, 3B, 4F**). It has been shown that structural heterogeneity introduces a new bias into nematode movement, which plays an important role in complex environments [51]. A monolayer of sand grains on agar surface has been applied before to introduce structural heterogeneity in *C. elegans’* environment and test its impact on worm trails [52]. Peb2les are heterogeneous (**Fig. 1**), and therefore can be used to explore the environmental, physical, and mechanical parameters that contribute to nematodes’ locomotory responses, and extract ethological insight. Peb2les can be combined with chemical cues, either administered in the NGM substrate or diluted in the particle solution, and they are compatible with microscopy and imaging techniques. These properties allow for ample experimental flexibility. Lastly, since Peb2les do not require sophisticated methods of fabrication, their potential for broad usage by the community is high.

### *C. elegans* locomotory behaviour in Peb2les

Nematodes in confined wet granular media [17, 53] were found to swim faster than in plain liquid. In our experiments, nematodes crawl on a moist agar substrate, layered with only slightly wetted particles, and their travelling speed depends on the density of the particle monolayer around the head and neck area (**Fig. 3A, S2**) but not on the density around the mid or posterior body (**Fig. S2**). Therefore, locomotory speed during crawling is adjusted based on density-dependent feedback as the worm heads into a new area and is not determined by input received elsewhere along the body. Despite the slight impediment, polystyrene granules of sizes analogous to the nematode’s body diameter do not block forward motion (**Fig. 3D**, **S3**).

Nematode locomotion is severely hindered in glass Peb2les, as reflected in low travelling speed (**Fig**. **3A**), compressed tracked paths (**Fig. 3B**, vs **2D, 2E**) and low *r* index (**Fig. 3D, S3**). This suggests that attractive forces between glass beads are too strong for worms to break through particle clusters. Instead, *C. elegans* move on top of the particle layer, however locomotion is severely impeded there, too (**Supplementary Video V4-V5**). Glass particles have high surface energy [54], hence they tend to adhere to surfaces. The latter can include both the NGM surface and the body surface of nematodes themselves. Strong attraction and cluster formation may be accentuated by ambient humidity [55], which may apply in moisture-rich NGM plates. The inability of nematodes to break through glass clusters can elucidate the boundaries of *C. elegans* locomotory adaptability. Moreover, nematodes in glass arenas sporadically lift and wave their heads, in a motion that resembles nictation [42], even though the rest of the body is not lifted (**Supplementary Video V5**, 0:10-0:12). Given that nictation is associated with dispersal strategy under harsh conditions [42], this enhances the notion that a glass arena is not a nematode-friendly environment.

Wild type and *mec-4* nematodes have similar travelling speed in all densities of polystyrene particles (**Fig. 4A**), and similar frequency distribution of their *r* index (**Fig. 4B**, **Fig. S3**). Therefore, sensory input on the nematode’s granular surroundings that informs locomotory adaptability does not seem to be collected by gentle touch neurons. Our findings agree with recent studies which report that regulation of nematode locomotory behaviour in granular environments does not involve soft-touch sensory neurons [48]. Particle packings, especially dense ones, present crawling *C. elegans* with resistance forces. This is mirrored in the finding that nematodes move more slowly in the presence of particles than they do in their absence (**Fig. 3A, 4A**), and in that increasing particle density around the head and neck areas reduces travelling speed (**Fig. 3A, 4A,** and **Fig. S2**). It is possible that these resistance forces are sensed by either proprioceptive neurons with stretch- or tension-sensitive mechanoreceptors [56, 57] or by mechanical sensors in the body wall muscles of the anterior body, such as PEZO-1 [58], or by both.

### “Blanket crawling” behaviour depends on granular terrain properties

Results from burrowing assays have shown that nematodes navigating dense 3D environments with mechanical resistance are challenged in a manner distinct from standard locomotion assays [59]. Similarly, exposing nematodes to an enriched, granular Peb2le environment reveals behaviours that are not possible to observe during standard behavioural assays. In one of them, nematodes in medium and high-density areas of polystyrene Peb2les spend a substantial fraction of time crawling under the particle layer (**Fig. 4C, 4D**). We call this “blanket crawling”. This is consistent with the well-established *C. elegans* burrowing behaviour on NGM plates [60] or other material [59, 61], and with observations about nematodes that immerse themselves in thick bacterial lawns [62].

Time spent crawling under the particle layer increases with particle density (**Fig. 4C**, grey). Compared to rearranging densely packed particles, which would require larger forces applied by the nematode, blanket crawling may be less energy demanding. Blanket crawling is also detected in *mec-4* mutants (**Fig. 4C**, green), which suggests that it is not mediated by touch sense, at least at medium particle densities, where wild type and *mec-4* behaviour is similar. At high particle densities, *mec-4* blanket crawling has a lower mean compared to wild type animals (**Fig. 4C**), but the difference is statistically borderline non-significant. This may imply a contributing but not required role for touch sense neurons.

Blanket crawling is not observed in glass Peb2les, where locomotion is significantly obstructed (**Fig. 3A, 3B, 3D**). This suggests that not only attractive forces among glass beads are too strong for worms to break through the bead clusters, but that attraction between the NGM substrate and glass beads is too strong for nematodes to crawl at their interface. This can be attributed to glass beads’ high hydrophilicity, with a lower nominal contact angle (∼45°) than that of polystyrene’s (∼90°), making particle-particle, particle-worm, and particle-substrate attraction strong [54]. This finding can provide insight to the physical properties of habitats that nematodes favour or avoid in the wild.

### Decision to enter new granular terrain may be linked to sensory sampling

Sensing the particles through touch affects nematodes’ initial decision to enter a new granular area (**Fig. 4E**), as shown in the case of diamond particles. In addition, failed attempts are often accompanied by a reversal (**Supplementary Video V5**), suggesting that the first few contacts with particles elicit a behaviour that resembles an escape reaction [63]. However, the fact that nematodes enter a Peb2le arena even if it takes several attempts, implies that the initial interaction does not discourage them from eventually exploring the new terrain. This touch-and-test sequence of attempts presents similarities with a recently reported “accept-reject” exploratory behaviour of *C. elegans* [64], in which animals sample a bacterial lawn they encounter for the first time, before they decide to move forward.

### Nematodes returning into granular terrain reveal touch-seeking behaviour

In contrast to other granular assays for nematodes [15, 17, 53, 65], Peb2les constitute an unconfined terrain, which animals can exit and revisit *ad libitum*. We found that most wild type nematodes return to polystyrene Peb2les within 40 sec of their initial exit (**Fig. 4G**, grey) and the returning behaviour cannot be attributed to chemosensory-driven attraction (**Fig. 5A**). The percentage of exits that results in returns are significantly reduced in *mec-4* mutants (**Fig. 4G**, green), suggesting that returning is driven, at least in part, by functional touch sense.

Mechanosensation is well studied in *C. elegans* [20–22, 66]. Even light touch at the anterior or posterior body evokes an avoidance response [67, 68], as nematodes perceive localized touch primarily as an aversive stimulus [21]. In polystyrene Peb2le arenas, a nematode’s entire body is in touch with particles (and the NGM substrate) almost continuously, regardless of whether the worm pushes through the particles or whether it crawls under them.

Nematodes show increased occupancy in Sephadex gel bead lawns [47], similarly to polystyrene microparticles that were used as comparison in the same study. Moreover, the mechanosensation of bacterial lawns triggers a mechanism that allows nematodes to remain in the food-rich environment [45]. In agreement with these findings, nematodes that have experienced the tactile-rich environment of a polystyrene Peb2le stay within the arena even after several hours (**Fig. 5B**). The tactile input received by *C. elegans* in Sephadex layers and in bacterial lawns is described as rewarding [47]; the finding that the majority of nematodes stay in the Peb2le arena or return to it could be explained by a similarly rewarding, touch-evoked, body-wide signal.

The ratio of returns that are realized via reversals or via other types of turns is maintained the same in both wild type and *mec-4* mutants (**Fig. 4H**). Possibly, motor circuits that do not require the contribution of mec-4-expressing neurons mediate the returning behaviour. Active touch-seeking in Peb2les may be related to *C. elegans* entrainment along physical wall-like barriers [69]. The underpinning mechanisms of the two behaviours could share common neuronal paths, since *mec-4* and *mec-10* do not exhibit entrainment [69], and *mec-4* display reduced touch-seeking behaviour in Peb2les (**Fig. 4G**).

In mammals, mechanosensory cues are linked to several forms of rewarding touch, like caressing [70] or even cuddling soft cloth for contact comfort [71]. To our knowledge, this is the first time that a behaviour resembling active touch-seeking is identified in *C. elegans*. Dissecting the underlying circuit would elucidate another nuanced behavioural expression of this nematode’s remarkable repertoire and could shed light on the evolutionary origins of this behaviour.

### *C. elegans* dynamically reshape the granular terrain

Crawling nematodes relocated neighbouring particles, and the worm-induced velocity of a particle (v_p_) declined exponentially with its distance (l_p_) from the moving worm body (**Fig. 6A, 6B**). This implies that dv_p_/dl_p_∼v_p_, meaning that the rate in which v_p_ decays with distance is proportional to v_p_ itself. This is typically a result of dissipative forces such as particle-particle friction and particle-substrate friction, which screen the worm-induced perturbance from far-field regions. There was no difference in particle displacement caused by wild type or *mec-4* (**Fig. 6B**), suggesting that the magnitude of forces exerted by moving nematodes was not determined by tactile feedback, but possibly by the mechanics of the motion itself.

Meanwhile, the exponential function form allowed the extraction of a single decay length (l_O_), marking a characteristic influence zone. In this case, the head-induced influence zone had an l_O_ that was twice as large as that of the full body (**Fig. 6B**). This may be a combined result of (i) the head-first locomotion, requiring the head to “spearhead” and penetrate into tightly packed particles while the rest of body simply follows through without having to displace particles as much, and (ii) the head performing large side-to-side sweep motion (**Fig. 2D, 2E**), which may relocate particles further.

Although the analysis of Delaunay triangles (**Fig. 6C**) does not distinguish the packing structure near or far from the worm in a statistically significant way, long-term changes in the landscape of Peb2les are clearly visible (**Fig. 7A, 7B, 7C**). Hexagonally ordered packing becomes more pronounced over time (**Fig. 7E**), with each particle having six neighbours in close contact. It would be far more difficult for the worms to displace these particles, as there is no room for compression except for them to be stacked, which would require stronger forces. The void areas that worms created (**Fig. 7B, 7D**) would be easier for them to navigate, in case of a revisit. In a way, the reshaped landscape of Peb2les would likely bias a worm’s future navigational choices.

Combined, these findings highlight the dynamic character of the Peb2les, and the ability of nematodes to reshape their environment in a way that depends on the surrounding medium physical properties. Because of that, Peb2les can be used to characterize body-environment interactions of nematodes that are models of neuromuscular disorders [60] and elucidate further the biomechanics of legless locomotion [72, 73].

## Conclusions

Peb2les are dynamic, versatile, and sensory enriched granular arenas, that can be successfully used to determine the limits of nematode locomotory adaptability. Our approach complements and advances prior findings, because Peb2les allow the expression of a broad spectrum of behaviours that are affected by environmental cues, and provide insight into behaviours that cannot be studied on standard culture plates. Our findings reveal that *C. elegans* display nuanced behavioural plasticity in granular terrains, in a way that depends on particle properties, like packing density, attraction forces, and surface texture of granules. We report previously uncharacterized nematode behaviours, like “blanket crawling” and active touch-seeking. Touch sense informs initial exploratory decision making and touch-seeking, but not locomotory adaptation. Moving nematodes reshape their granular environment in a continuous and dynamic manner, revealing ecologically relevant preferences and trends.

## Acknowledgements

We thank Zhaochen Yang for help with video analysis; members of the Gourgou Lab Jonathon McSwain and Kareem Hassouna for help with nematode cultures, and Bianca Pereira and Jonathon McSwain for comments on the manuscript; and Achilleas Anastasopoulos for assistance with data processing. Nematode strains were provided by the CGC, which is funded by NIH Office of Research Infrastructure Programs (P40 OD010440).

## Authors’ contributions

HX: experimental design, data analysis, code writing, manuscript writing, manuscript review. SM: data collection, data analysis, help with manuscript writing. YP: data analysis and code writing. AS: data analysis and code writing. EG: conceptualization, experimental design, data collection, data analysis, research supervision, manuscript writing, manuscript review, funding acquisition.

## Data availability

Data related to this work are provided in the Appendix, Supplementary Figures, and Supplementary Videos. More videos will be available at an accessible repository upon acceptance of the manuscript.

## Supplementary Information

## Appendix

## Supplementary Videos

**Supplementary Table 1:**
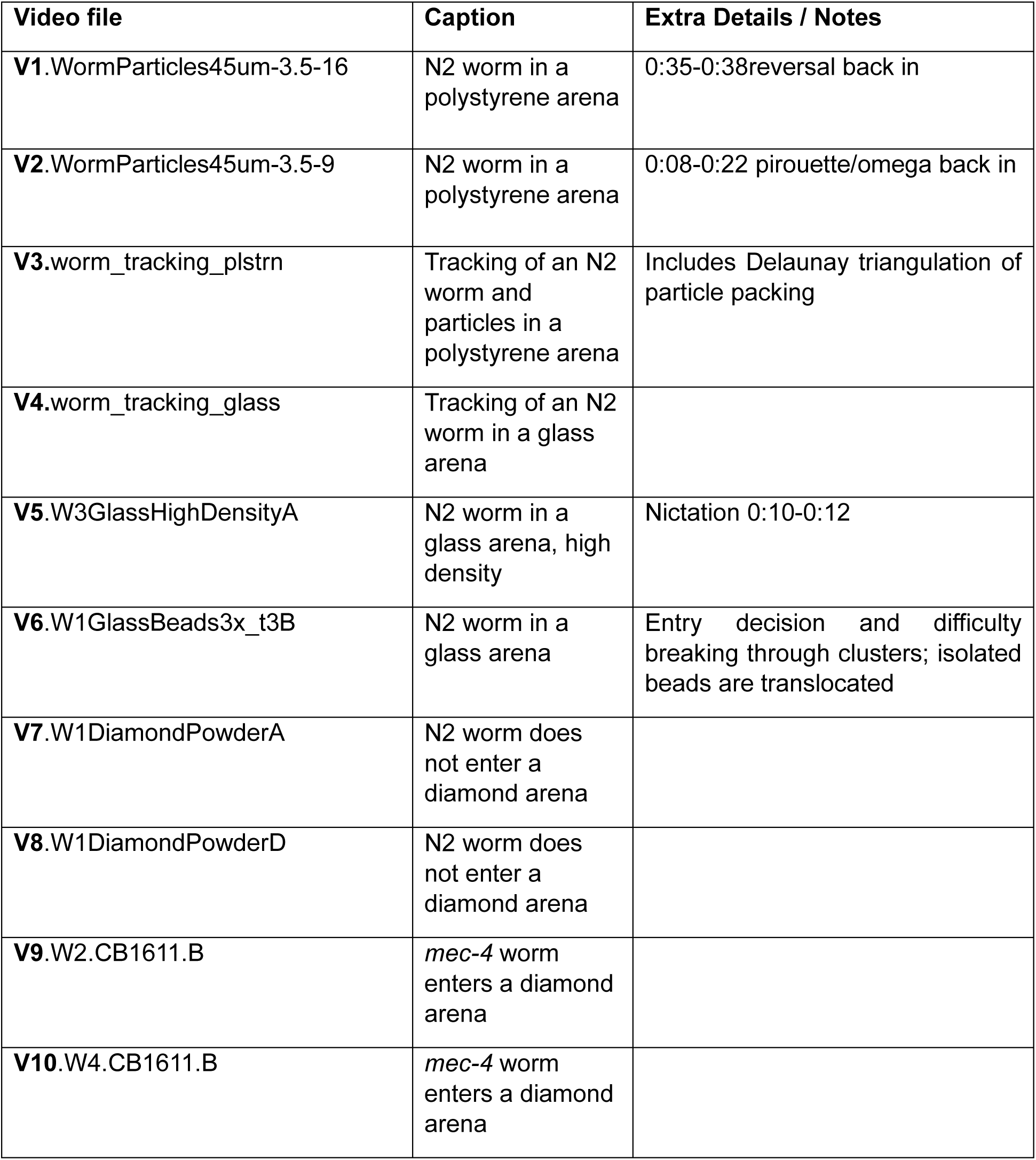
List of supplementary videos, with captions and details.

| Video file | Caption | Extra Details / Notes |
| --- | --- | --- |
| **V1.**WormParticles45um-3.5-16 | N2 worm in a polystyrene arena | 0:35-0:38reversal back in |
| **V2.**WormParticles45um-3.5-9 | N2 worm in a polystyrene arena | 0:08-0:22 pirouette/omega back in |
| **V3.**worm_tracking_plstrn | Tracking of an N2 worm and particles in a polystyrene arena | Includes Delaunay triangulation of particle packing |
| **V4.**worm_tracking_glass | Tracking of an N2 worm in a glass arena |  |
| **V5.**W3GlassHighDensityA | N2 worm in a glass arena, high density | Nictation 0:10-0:12 |
| **V6.**W1GlassBeads3x_t3B | N2 worm in a glass arena | Entry decision and difficulty breaking through clusters; isolated beads are translocated |
| **V7.**W1DiamondPowderA | N2 worm does not enter a diamond arena |  |
| **V8.**W1DiamondPowderD | N2 worm does not enter a diamond arena |  |
| **V9.**W2.CB1611.B | *mec-4* worm enters a diamond arena |  |
| **V10.**W4.CB1611.B | *mec-4* worm enters a diamond arena |  |

## Supplementary Figures

**Figure S1:**
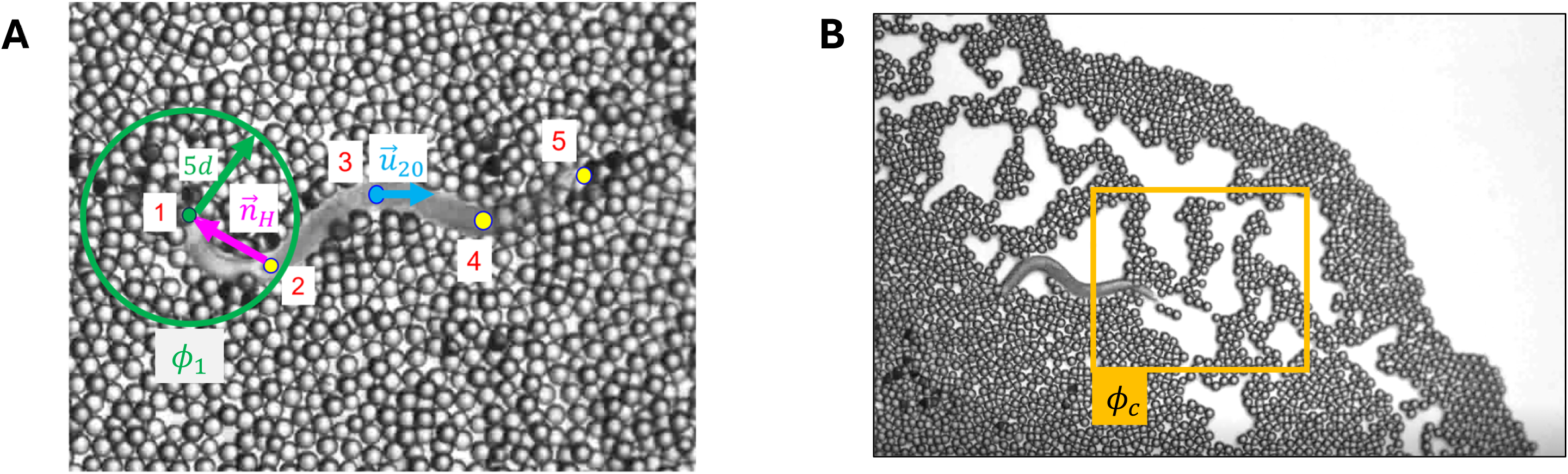
Particle packing fraction calculations and vectors for direction index calculations. (A) For each identified worm body node (numbered as 1-5, head to tail, see also Fig. 2C), we calculate the packing fraction by sampling all the pixels within 5 particle diameter (5d, ∼0.22 mm) away from each node. An example is shown for the head node, *i.e.,* ¢_1_ particle density (green circle). u_20_: displacement vector over 20 experimental frames; n_H_: neck to head vector. The length of u_20_ is typically about 1/5 worm body length, which is a large enough value comparing to the tracking error for the centre node of the worm. See also Figs. 3A, 3C, 4A. (B) Area in front of the worm, used to calculate particle density ¢_c_ for crawling-under-the-particles behavior (*i.e.,* blanket crawling, see Fig. 4C), square dimensions: 1 mm x 1 mm.

**Figure S2:**
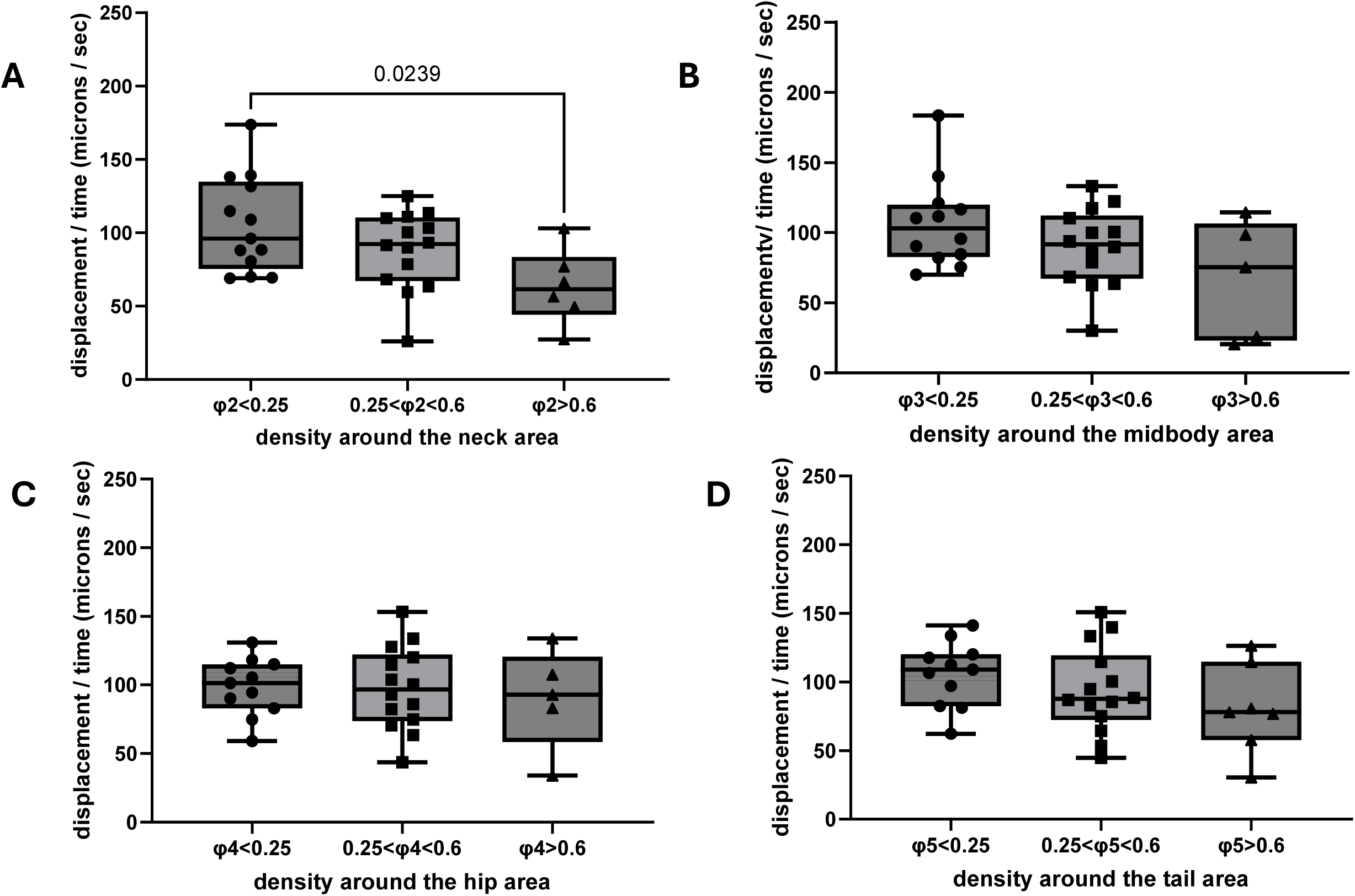
Locomotory performance of *C. elegans* with respect to particle density around different body areas. Displacement per second, of N2 wild type nematodes, in polystyrene Peb2le arenas, with increasing particle density around the neck (**A**), midbody (**B**), hip (**C**), and tail (**D**) areas (see also Fig. 2C and Fig. 3C for position of body nodes). Each dot represents displacement of one worm, n = 13 worms processed. The number of worms that visited areas of different densities varied. Horizontal lines: mean values, boxes: the central 50% of data, bars: min and max values. One-way ANOVA followed by Tukey’s multiple comparisons test; only p-values <0.05 are shown. For displacement with respect to particle density around the head area, see Fig. 3A.

**Figure S3:**
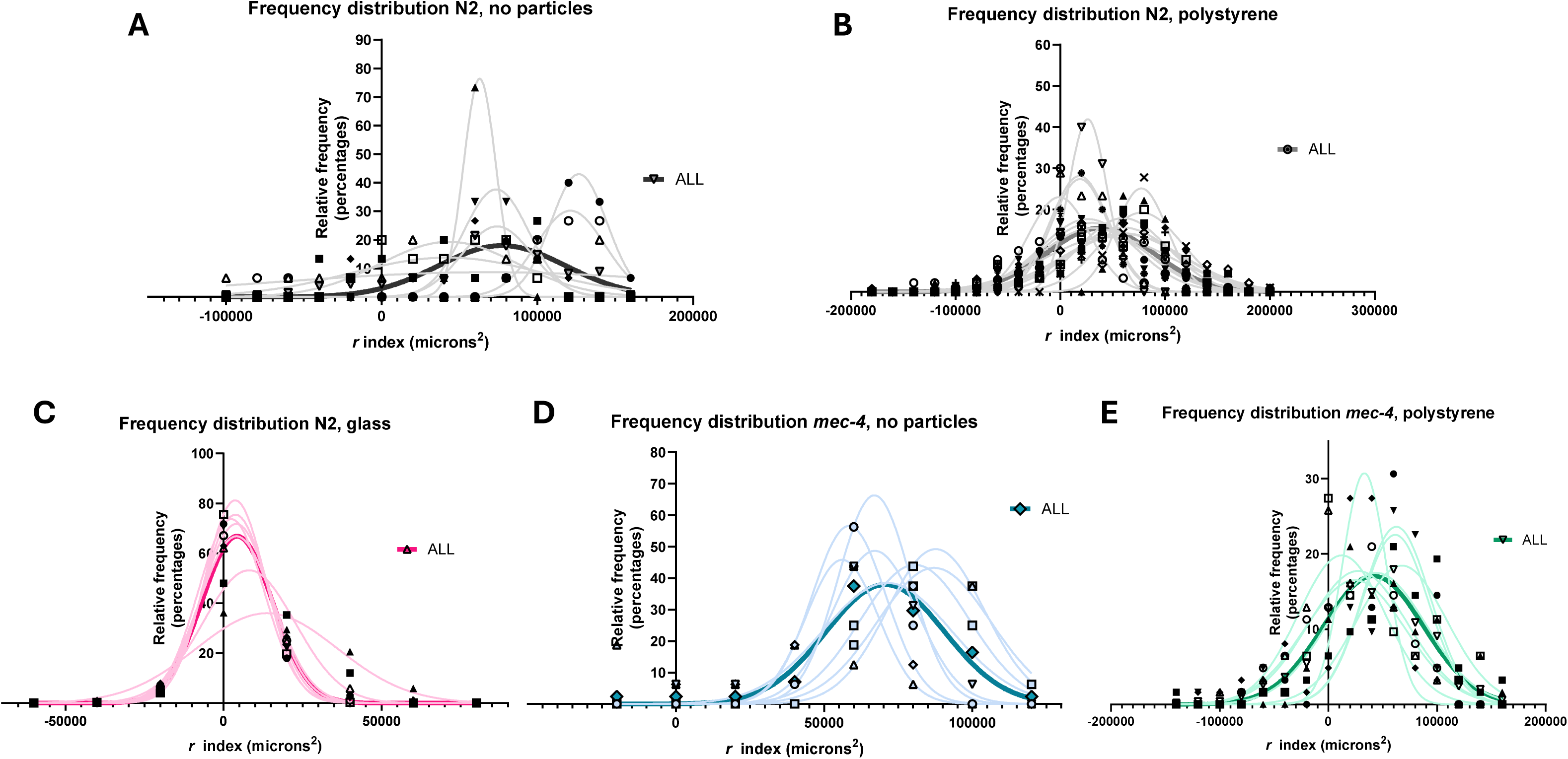
Direction index *r* for all nematodes analyzed, per arena type and strain. Relative frequencies (percentages) distribution of direction index *r* for the assessment of backward/forward motion (see Methods and Appendix) of N2 wild type in the absence of particles (**A**) and in polystyrene (**B**) and glass (**C**) Peb2les, as well as of *mec-4* in the absence of particles (**D**) and in polystyrene Peb2les (**E**). Individual shapes indicate bin centre of histogram for frequency distribution; curves represent nonlinear fit for Gaussian distribution. *r*>0 indicates forward motion, *r*<0 indicates backward motion. Darker curves, labelled as “ALL”, indicate the mean distribution for all nematodes of the group, also displayed in Figs. 3D and 4B.

**Figure S4:**
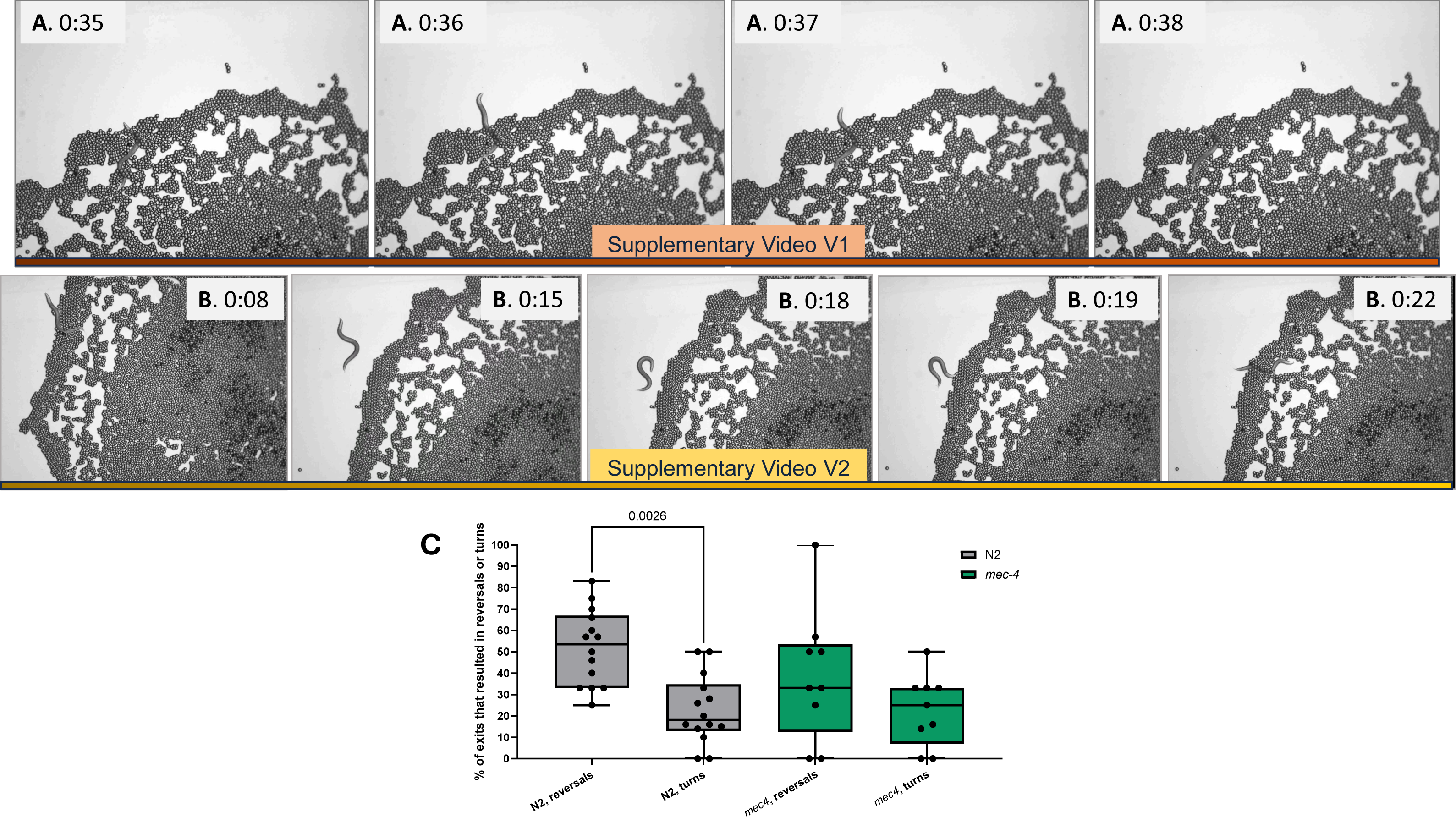
Touch-seeking behaviour in Peb2le arenas, and the role of *mec-4*. **(A panels, top row)** Sequence of snapshots of a return via a reversal, see Supplementary Video V1 and indicated time points in seconds. **(B panels, middle row)** Sequence of snapshots of a return via a sharp (omega) turn, see Supplementary Video V2 and indicated time points in seconds. (C) % of exits that were realized by either a reversal or a turn (for events definitions see Methods), for N2 wild type (grey) and *mec-4(e1611)* (green) nematodes. See also Fig. 4H. Wilcoxon matched-pairs signed rank test. Only p-values <0.05 included. Horizontal lines: mean values, boxes: the central 50% of data, bars: min and max values.

**Figure S5:**
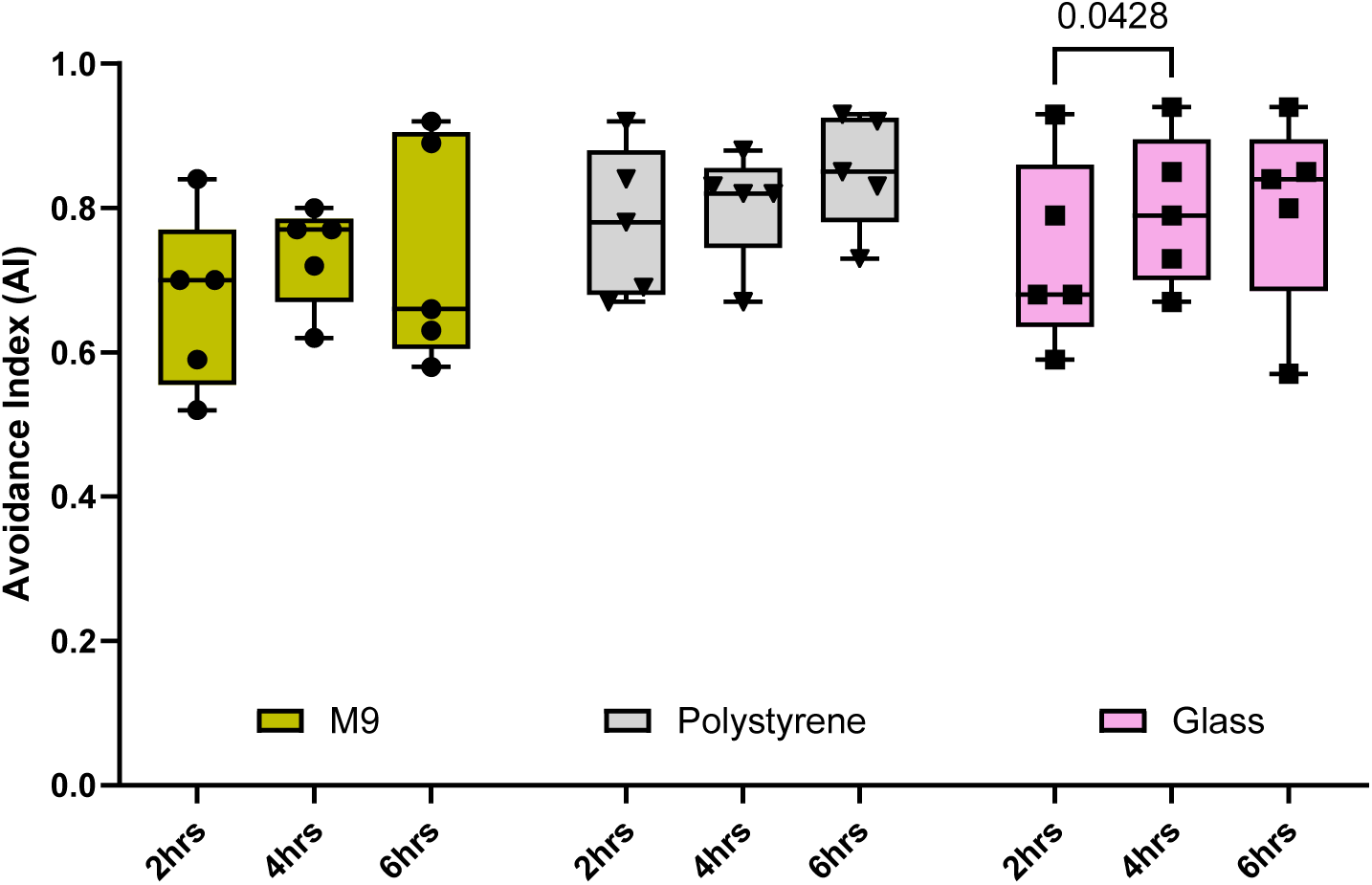
Long term avoidance of Peb2le arenas. Avoidance index at different time points (2 hr, 4 hr, 6 hr) for N2 wild type nematodes and particles used in Peb2le arenas. Particles were mixed with M9 buffer. Two-way ANOVA, followed by Tukey’s multiple comparisons test. Only p-values<0.05 are shown. Comparisons against plain M9, no particles, and among groups of the same particle type.

