## Appendix for "Behavioural plasticity in *Caenorhabditis elegans* navigating dynamic granular environments"

#### Nematode tracking via Deep Learning

The methodology of worm tracking was twofold (**Fig. 2A**). The first step was a semantic image segmentation to identify all pixels in an image that belonged to the worm. Specifically, we employed the DeepLabV3+ architecture with ResNet-34 as the encoder [1, 2], which is effective at handling multi-scale contextual information, *e.g.*, small-scale tail tip and large-scale curvature of the undulating worm body, and can yield sharp object boundaries. This is a supervised learning algorithm for which the input was the grey scale raw image (**Fig. 2A**, left panels) and the label was a masked image (**Fig. 2A**, right panels), in which pixels belonging only to the worm were highlighted. The masks for training were created manually using the brush tool in Adobe Photoshop. We employed the binary cross entropy with logits loss (BCEWithLogitsLoss in PyTorch) as the loss function and trained the neural network over 30 epochs with the Adam optimizer [3], with a learning rate of 0.001. An example prediction for an independent testing image is provided in **Fig. 2C**, in which the majority of pixels belonging to the worm are correctly identified (lavender blue).

The second step for worm tracking was post-processing, to extract worm information from each image prediction and track the worm over a time series of predictions. From the segmented image, we identified the largest contiguous area (*i.e.*, the worm) and applied a morphological closing operation to smooth the boundary and ensure topological continuity. The refined area was then reduced to its centreline via skeletonization and was combined with a path finding approach to sort the skeletonized pixels from one end of the worm body to the other. Next, we further reduced the centreline to its five body nodes, which are at $0^{th}$, $25^{th}$, $50^{th}$, $75^{th}$, and $100^{th}$ percentiles of its total length, representing the head, neck, midbody, hip, and tail (**Fig. 2C**). These points were tracked over time to obtain the trajectory of the worm. At the beginning of each video, we manually identified the node corresponding to the head (**Fig. 2C,** green), which was then applied throughout a time trajectory. For displacement over time, as shown in **Fig. 3** and **Fig. 4**, we calculated displacement (in pixels) of the midbody node over 20 frames (frame rate 13.74 frames per second). Then we converted to microns per second, as shown in **Figs. 3** and **4**.

While **Fig. 2** and the corresponding supplementary videos demonstrate success in most common scenarios, it is possible that in an image with densely packed particles, the tip of the head and especially of the tail are not identified, resulting in the overall identified length of the worm to be shorter. This does not significantly impact the midbody node of the worm, and we use this node for worm displacement analyses. In rare instances, the worm curls up, *i.e.,* as in an omega turn, making tracking difficult. These frames (<3%) were disregarded.

### Particle tracking via Deep Learning

For particle identification, we applied the same semantic segmentation methodology, but instead used an existing package, Bellybutton [4], for identifying all pixels that belonged to the particles, including stacked particles and particles overlapping with the worm (**Fig. 2A, 2B**). Bellybutton requires similar input and labelling as the worm tracking method, but with the particles highlighted (**Fig. 2B**). An example prediction is provided in **Fig. 2C**, with the identified pixels highlighted in pink. For non-stacking particles, we proceeded with using the circular Hough transform to identify the centroids, based on which we quantify their motion and packing structure, see section "Particle kinematics and rearrangement by traveling nematodes".

### Calculation of particle density

For each identified worm body node 1-5 (**Fig. 2** and **Supplementary Figure S2A**), we calculated the packing fraction by sampling all the pixels within the radius of 5 particle diameters from the node, with the estimated diameter $d = 20$ pixels. Then, for each node $i$, we calculated the packing fraction as

$$\phi_i = \frac{number\ of\ pixels\ that\ belong\ to\ the\ particles}{all\ pixels}$$

Note that the pixels corresponding to the worm are also included in the denominator.

### Calculation of direction index $r$ for the estimation of backward/forward motion

To determine whether a worm moves forward or backward, we utilized the worm's displacement vector, $\vec{u}_{\text{m}}(\Delta t)$, measured at the midbody node over 50 frames, *i.e.,* time interval $\Delta t = 3.65$ s, and the vector pointing from the worm's neck to head, $\vec{u}_{\text{nh}}$ at the starting instance of the interval (**Fig. 3C**). Their internal product defines a direction index, $r = \vec{u}_{\text{m}} \cdot \vec{u}_{\text{nh}}$, with $r > 0$ indicating forward motion and $r < 0$ indicating backward motion. When the worm displays little movement over $\Delta t$ or when the head orientation is perpendicular to the body displacement, $r \approx 0$. We visualized the results by plotting the relative frequency distribution of $r$ (**Fig. 3D**, **4B**, and **Supplementary Figure S3**). To avoid skewed data due to the unequal number of data points for each video/worm/condition, we randomly sampled data sets at a final number of data points equal to the least number of data points of each set (MATLAB 2023a, MathWorks). We checked the robustness of sampling by plotting and comparing the mean, standard deviation, and median of the unsampled and sampled data.

### Chemotaxis assay

The chemotaxis assay was performed according to a well-established four-quadrant protocol [5]. Polystyrene 45.0 µm particles come as 2.7% aqueous suspension by the manufacturer (Polysciences, USA). Initially, 70 µL were precipitated by centrifugation at 4300 rpm for 30 s, and after the supernatant was discarded, an equal volume (~10 µL) of sterile M9 was added in the pellet and mixed. Glass beads (Cospheric, USA) and diamond powder (Pureon, USA) come in solid form; 10 mg of glass beads or 75 mg of diamond powder were suspended in 50 μL sterile M9. For each particle type, 5 μL of particle suspension was pipetted at each test quadrant and was left to dry. For the positive control experiments, 5 μL of 0.5% (w/v) diacetyl solution (TCI, USA), instead of the particle suspension, was applied at the test quadrants. 5 μL of M9 buffer was pipetted at the centre of the control quadrants for all experiments. Next, 5 μL of 0.5 M sodium azide was added at the centre of all quadrants and was allowed to dry. All solutions were placed at equal distance of 2 cm from the plate centre.

A synchronized population of Day 1 adult hermaphrodites was obtained, following standard process [6] with slight modifications [5]. Nematodes were collected using sterile M9 and were centrifuged at 6600 rpm for 10 sec. The supernatant was removed, and worms were washed with 1 mL fresh M9. This washing step was performed three times. After final centrifugation, ~100 μL of M9+worm suspension was retained in each tube. A 5 μL aliquot of worm solution was pipetted onto an unseeded NGM plate to estimate worm numbers, and an appropriate volume, containing ~50-150 worms, was used per chemotaxis plate.

The worm+M9 suspension was introduced at the plate centre. After excess liquid was absorbed (~5 min), assay plates were sealed with parafilm and incubated for 1 h in 20°C. Next, assay plates were chilled at 4°C for 15 min, worms in each quadrant were counted, and the chemotaxis index was calculated, as *Chemotaxis Index* = ($N_{test}$ - $N_{control}$) / ($N_{total}$), where $N_{test}$ is the number of worms in both test quadrants, $N_{control}$ is the number of worms in both control quadrants, and $N_{total}$ is the number of total scored worms. Five to eight replicates for each condition were performed for each assay, across three different days.

### Avoidance assay

For the avoidance assay, a previously published protocol was followed [7]. A synchronized population of Day 1 adult hermaphrodites was obtained, following standard process [6]. On the day of the assay, the particle solution, in M9 or in OP50, was prepared freshly. For glass and diamond particles, we followed the same process as for the chemotaxis assay (see above). For the polystyrene particles, 250 μL of manufacturer's solution were centrifuged at 4300 rpm for 30 sec, and the supernatant was discarded. Next, 70 μL of sterile M9 was added in the pellet, mixed thoroughly, and the mix was pipetted at the centre of the assay plate, to form a monolayer with a circle of ~1.7 cm in diameter. For the control experiments, 50 μL of M9 or OP50 were used, instead of a particle solution (**Fig. 5** and **Fig. S5**). Assay plates were left to air dry. Day1 worms were collected and washed with M9, and ~30-60 worms suspended in ~5 μL of M9 were transferred onto the particle/bacterial lawn. The plate was sealed with parafilm and incubated at 20°C. Avoidance was scored 2, 4 and 6 hr later. Results were quantified by calculating the fraction of worms staying on the particle/bacterial lawn over the total number of worms in the assay plate, as *Avoidance Index* = $N_{off}/N_{total}$, where $N_{off}$ is the number of worms outside the lawn and $N_{total}$ is the number of total worms on the plate. Five replicates for each condition were performed for each assay, across three different days.

### Particle kinematics and rearrangement by traveling nematodes

We calculated the average particle velocity ($v_p$) as a function of each particle's distance, $l_p$, from the worm [8], the latter defined in two ways. First, we sorted the particles according to the shortest distance from a particle's centre to the worm's surface contour, *i.e.*, the boundary of the highlighted worm body (**Fig. 6A**). Second, we sorted the particles according to their distance to the worm's head node. Both results are shown in **Fig. 6B** for N2 wild type and *mec-4* mutants. For both definitions of $l_p$, the particle motion decays rapidly with increasing $l_p$, and particles closer to the head display larger worm-induced motion than particles elsewhere. Results for the two worm strains are identical. The inset of **Fig. 6B** shows the same result on a semi-log scale, indicating that the particle motion decay is exponential, *i.e.*, $v_p \sim e^{-l_p/l_0}$, with the fitted decay length $l_0 = 33$ μm for the body-based calculation and $l_0 = 71$ μm for the head-based calculation (note that the particle diameter is 45 $\mu\mathrm{m}$).

To determine how a crawling nematode rearranges the particles nearby, we characterize the packing structure of the particles using Delaunay triangulation (**Fig. 6A**). We partitioned them into near-field (yellow) and far-field (blue) triangles, while discarding triangles that overlaps with the worm's body (black). We examined the average triangle area, $A_{\mathrm{tri}}$, as a function of particle packing density $\phi$, which is averaged over $\phi_1 - \phi_5$ (**Fig. 6C**). The difference between near- and far-field triangles was found to be statistically insignificant at all particle densities (p-values >0.05). Note that for both the kinematics and the structure analysis we did not consider cases with $\phi > 0.6$, as in those cases the worm moves mostly under the particles (**Fig. 4C, 4D**) and tracking of particle centres becomes unreliable due to occasional particle stacking.

#### Particle packing and Fast Fourier Transform

To detect nematode-induced long term modifications of the Peb2le arena we compared images of the entire arena taken before the worms are introduced and 3 h after their introduction (**Fig. 7A, 7B**). We calculated the difference between the two images, with the intensity difference shown in **Fig. 7C**. To further quantify how the arena is changed, we sampled sub-regions of the two raw images (**Fig. 7A, 7B**) with a window size of $0.71 \times 0.71\ \mathrm{mm}^2$, in which significant nematode-induced differences were detected from **Fig. 7C**. For each sub-region, we applied a two-dimensional Fast Fourier Transform (FFT, **Fig. 7D**, inset). Finally, we performed a radial average of the FFT result, further average over all sub-regions for each image, and obtained a relation between the FFT intensity, $|\tilde{I}(f)|^2$, and the spatial frequency, $f$, for each image (**Fig. 7D**). Fourier analysis was performed in MatLab (MathWorks, USA).

The Fourier transform-based quantification of the sub-domains with significant packing structure difference (**Fig. 7E**) shows a more pronounced peak at $1/d = 0.022\ \mu\mathrm{m}^{-1}$in the "after" image than the "before" image, indicating tighter packing between particles. At lower wavenumber, $f < 0.02\ \mu\mathrm{m}^{-1}$, the measured FFT intensity $|\tilde{I}(f)|^2$ is also higher in the "after" image, which should correspond to larger-scale white spaces in the image, *i.e.*, the void areas created by the worms.
